# Computational modeling of AMPK and mTOR crosstalk in glutamatergic synapse calcium signaling

**DOI:** 10.1101/2022.08.17.504291

**Authors:** A. Leung, P. Rangamani

## Abstract

Neuronal energy consumption is vital for information processing and memory formation in synapses. The brain consists of just 2% of the human body’s mass, but consumes almost 20% of the body’s energy budget. Most of this energy is attributed to active transport in ion signaling, with calcium being the canonical second messenger of synaptic transmission. Here, we develop a computational model of synaptic signaling resulting in the activation of two protein kinases critical in metabolic regulation and cell fate, AMP-Activated protein kinase (AMPK) and mammalian target of rapamycin (mTOR) and investigate the effect of glutamate stimulus frequency on their dynamics. Our model predicts that frequencies of glutamate stimulus over 10 Hz perturb AMPK and mTOR oscillations at higher magnitudes by up to 70% and area under curve (AUC) by 10%. This dynamic difference in AMPK and mTOR activation trajectories potentially differentiates high frequency stimulus bursts from basal neuronal signaling leading to a downstream change in synaptic plasticity. Further, we also investigate the crosstalk between insulin receptor and calcium signaling on AMPK and mTOR activation and predict that the pathways demonstrate multistability dependent on strength of insulin signaling and metabolic consumption rate. Our predictions have implications for improving our understanding of neuronal metabolism, synaptic pruning, and synaptic plasticity.

**Key Points:**

- Neurons consume disproportionate amounts of cellular energy relative to their mass, indicating the importance of energy regulation in information processing in the brain.
- AMP activated protein kinase (AMPK) is thought to be the biochemical link between energy consumption in neuronal information processing and synaptic plasticity.
- Computational model investigating the crosstalk between high-frequency glutamatergic calcium signaling and AMPK activation in neurons predicts multistability in AMPK and mammalian target of rapamycin (mTOR) activation.
- Our models predict a frequency-dependent response in AMPK and mTOR activation that also scales according to insulin signaling and energy consumption. The oscillatory behavior depends on both intracellular and extracellular factors, such as energy consumption and insulin signaling.
- This work elucidates the role of insulin and insulin resistance in regulating neuronal activity, through computational modeling the metabolic response of energy stress resulting from calcium signaling.

## Introduction

Calcium signal transduction in dendritic spines during neuronal signaling is closely linked to synaptic plasticity [1–5]. Synaptic plasticity is the structural and molecular modification of synapses resulting in sustained changes in synaptic signaling strength [2,6,7]. There are many potential mechanisms by which calcium signaling leads to long-term plasticity (LTP), the process by which synaptic connections are strengthened with high-frequency stimulus, but the relative contribution of these different mechanisms is unclear [8]. One such mechanism is metabolic plasticity, which is the adaptation of cellular energy production in response to metabolic stress associated with calcium signaling [9–11] and is the focus of our current work. Electrical signaling, including the reversal of presynaptic and postsynaptic ion fluxes, accounts for a vast majority of ATP consumption in mammalian neuron signaling [12,13]. During an action potential, energy consumption is estimated to transiently increase up to 130 percent over basal ATP flux with most of this energy attributed to ion signaling, actin and microtubule turnover, and lipid and protein translation [12–14]. Therefore, the coupling of metabolic plasticity with ion dynamics is necessary for dendritic spines due to a drastic increase in energy consumption during neuronal signaling [11]. In addition, there are a variety of energetically expensive processes in the dendritic spine that are necessary to induce long-term potentiation (LTP) including actin remodeling, protein translation, endocytosis, and exocytosis [14]. The frequency of neurotransmitter signaling (Figure 1a) is believed to be critical in the induction of LTP in dendritic spines and also increases the consumption of cellular energy due to the increased rate of active transport of ions to restore resting potentials [13].

**Figure 1:**
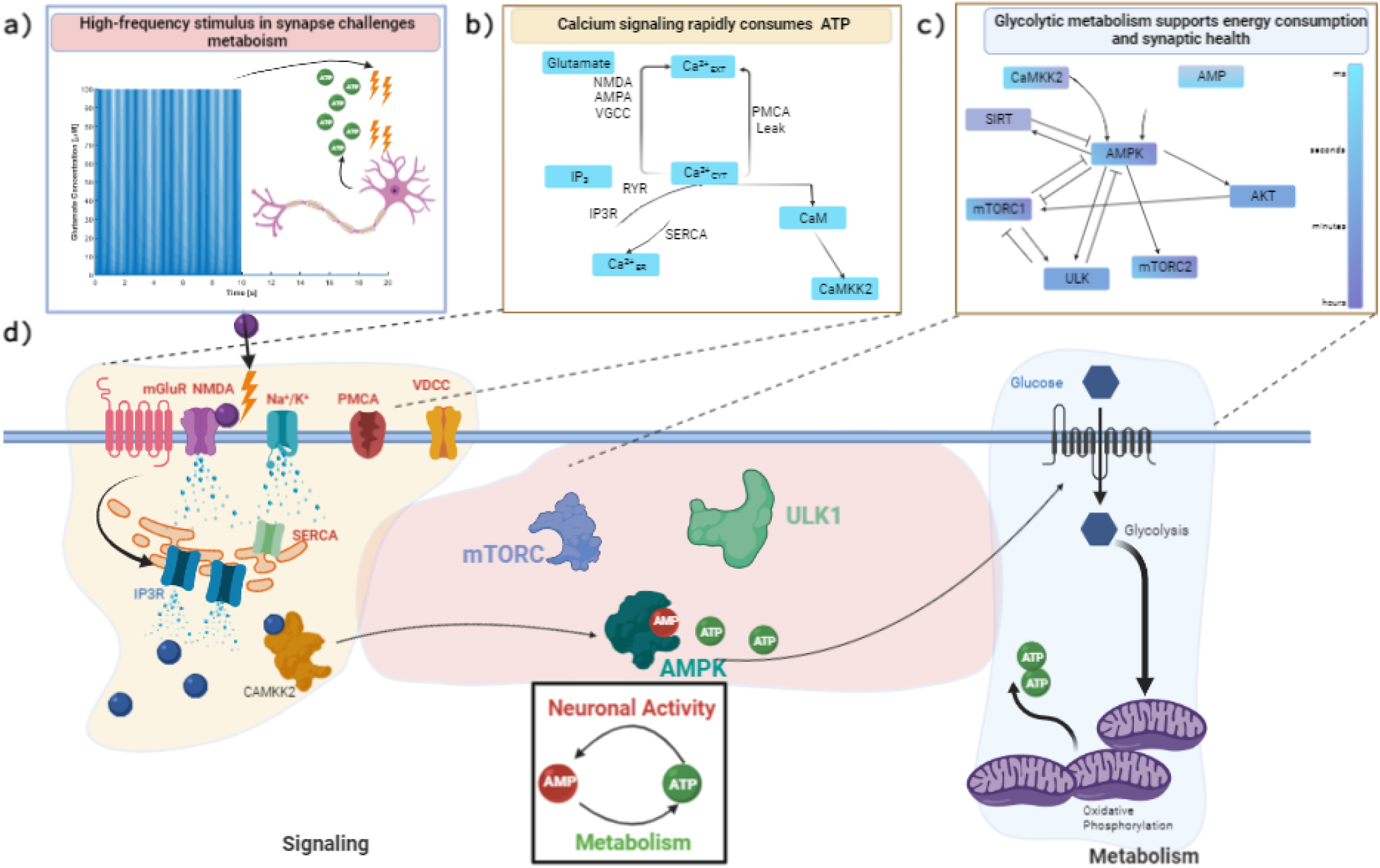
Synaptic signaling consumes energy to transduce neuronal signals and support neuronal function. During high-frequency synaptic signaling, represented with glutamate stimulus in **a)**, proportionally larger quantities of ATP are allocated to restoring resting potential of ions, like those shown for calcium illustrated in **b)**. At high frequencies, this may challenge the energy production capacity of neurons, which utilize glycolysis and oxidative phosphorylation to convert glucose to pyruvate and then produce ATP in dendritic mitochondria. The decreasing cellular energy state (higher AMP/ATP ratio) promotes the phosphorylation of AMPK, a kinase which promotes the production of cellular ATP, but also has an intricate feedback loop with mTORC1 and mTORC2 downstream of the insulin signaling cascade, shown in **c)**, which have implications in protein translation and synaptic plasticity. In this work, we develop and analyze a computational model to study the interactions of these pathways, illustrated in **d)**, in response to a synaptic stimulus across several timescales.

The two primary means of energy production within neurons are glycolysis and oxidative phosphorylation in the mitochondria [15,16]. Glycolysis, the metabolism of glucose in the cytosol, is often upregulated during neuronal stimulus through increased import of glucose from extracellular space and protein kinase-dependent activation of enzymes. [17,18]. Mitochondrial oxidative phosphorylation produces a large portion of cellular energy that supports increased energy demand. However, oxidative phosphorylation requires pyruvate and oxygen consumption to generate ATP in the mitochondrial matrix through mitochondrial metabolism of pyruvate driving the electron transport chain. In actively signaling neurons, mitochondria are observed to form spatially stable pools near the base of dendritic spine, suggesting that they can cater to the increased ATP demand by localizing ATP production proximal to the dendritic spine [19]. In conjunction, it has been observed that calcium influx from the extracellular space impacts the mitochondrial membrane potential, resulting in higher ATP generation [20]. This provides a route for calcium, the main second messenger system in signaling, to interact directly with the energy production in a frequency-dependent manner. The thermodynamic constraints of ATP production in the mitochondria are explored in [21].

Calcium influx (Figure 1b) indirectly modulates neuronal glycolysis and mitochondrial metabolism via enzymatic activation (CAMKII [22]), but the means by which neurons regulate the energy production local to the dendritic spine remains unexplored [20]. A possible avenue for feedback between neuronal energy consumption and metabolic production is through cellular energy sensors and kinases. Adenine monophosphate-activated Protein Kinase (AMPK) is a candidate molecule for the coupling between calcium dynamics and cellular metabolism due to its unique function as the cellular energy stress sensor [20,23]. The *γ* subunit of this heterotrimeric protein is able to adapt to changing ratios of AMP and ATP, exposing a phosphorylation site and enabling kinase functionality [24]. Upstream protein kinases of AMPK include CAMKK2, LKB1, AKT, mTOR, and PI3K, and may regulate AMPK activity depending on phosphorylation site [22,25,26]. The downstream targets of AMPK include phosphofructokinase (PFK), mitochondrial biogenesis, and inhibition of energy consuming pathways like lipid synthesis and gluconeogenesis. Calcium-Calmodulin activated Kinase Kinase 2 (CaMKK2) enables direct feedback from the calcium signaling pathway to AMPK. Calcium also creates an energy imbalance because of increased energy demand by ion channels, which increases AMP/ATP ratio and activates AMPK [11,22,23,27]. AMPK activated by calcium, phosphonucleotide binding, or allosteric activation pathways leads to feedback to dynamically enhance energy production. AMPK has been shown to increase the import of glucose into neurons by promoting trafficking of glucose import receptors GLUT3 to the plasma membrane [28]. Furthermore, AMPK upregulates glycolysis by modulating PFK activity during energy stress in cardiac cells as well as mitochondrial morphology in neuronal axons [29, 30].

One critical downstream target of AMPK is mammalian target of rapamycin (mTOR); select interactions are shown in Figure 1c. mTOR is a well-conserved protein across mammalian cell types, but occupies a unique niche in neurons. While classically mTOR forms a protein complex that regulates protein translation and mitophagy [31], mTOR has also been shown to be critical in synapse formation [11,31–33]. In presynaptic boutons, mTOR forms a complex with Rictor, mTORC2, which is necessary for bouton formation [31]. In postsynaptic dendritic spines, mTOR forms a complex with Raptor, mTORC1 [11]. mTORC1 has been shown to influence the translation and transport of α -Amino-3-hydroxy-5-methyl-4-isoxazolepropionic acid (AMPA) receptors and scaffolding proteins needed for the clustering of AMPAR, a critical step in synaptic plasticity [34]. Abolishment of mTOR activity in both presynaptic and postsynaptic cells has been performed in live neurons and shows a decrease in the number of spines and plastic ability [32, 34]. The exact mechanisms by which the calcium-AMPK-mTOR signaling axis is regulated in neurons are yet to be explored. In particular, exploration is needed to bridge the activity of calcium signaling, which typically occurs on the timescale of milliseconds, to the function of protein kinases (AMPK, mTOR, AKT), which can occur on the timescale of hours to days. AMPK and mTOR are also both downstream targets of the insulin receptor substrate (IRS) signaling cascade, which can bridge the intracellular response to a systems level change in insulin. Downstream of mTORC1 are several transcription regulators, for example, Sirutitin (SIRT1) has been shown to regulate neuroprotective pathways and may have a role in dendritic spine formation [35, 36]. A multi-timescale model that represents these different systems can shed light into the crosstalk between these two pathways.

In this work, we use computational modeling to explore how the calcium-AMPK-mTOR signaling axis could couple neuronal energy states and mTOR activity. Computational modeling has vastly contributed to our understanding of neuronal signaling and synaptic plasticity [1–4,37–39]. However, very few of these models feature the importance of metabolic feedback mechanisms known to be critical in synaptic formation and activity. We built our model based on prior models in the literature, utilizing a calcium signaling model that incorporates the effect of endoplasmic reticulum (ER) and mitochondria on calcium signaling and mitochondrial ATP production [40]. We complement this calcium model with a model linking cellular energy state via adenine nucleotide balance with AMPK activation [41]. Finally, we explore the AMPK and mTOR crosstalk, by integrating the insulin signaling model within a calcium and metabolism model [26].

Our model predicts that a wide range of stability behavior is possible depending on glutamate stimulus, neuronal energy consumption, and the behavior of downstream components of the insulin signaling pathway. Glutamate signaling frequency can influence key characteristics in the dynamics of vital protein kinases and can drastically increase the magnitude of AMPK and mTOR phosphorylation and total signaling observed during stimulus. Additionally, we observe that internal metabolism (ATP hydrolysis), external metabolism (insulin signaling), and neuronal frequency directly influence the system response, with parameter-dependent oscillations. The multiple stable states of this system implies that the crosstalk between extracellular signals (Figure 1d), such as gluatamate and insulin, culminate in changes to internal metabolic systems such as AMPK and mTOR activation rates. Further exploration of the crosstalk between pathways may elucidate the relation between neuronal metabolism and signal processing in dendrites.

## Methods

### Model Description

All differential equations are listed for Table 1 and the fluxes for each reaction are listed in Tables 2 to 4. Each parameter value used in the model is listed in Tables 5 to 8. Here, we describe the various modules and the original sources for the reactions, where applicable. The system is composed of models representing AMPK and mTOR activation (Table 2), calcium dynamics (Table 3), and receptor dynamics NMDAR and AMPAR (Table 4).

**Table 1:** Differential Equations used in the model.

| Index | Equation |
| --- | --- |
|  | Nucleotides and AMPK |
| 1 | $\frac{d}{dt}[ATP] = -r1 + r2 - AK - CK - 2 * J_{ATP,Ca} + rc,$ |
| 2 | $\frac{d}{dt}[ADP] = r1 - r2 + 2AK + CK + 2J_{ATP,Ca},$ |
| 3 | $\frac{d}{dt}[AMP] = -AK - r20 - rc,$ |
| 4 | $\frac{d}{dt}[PCr] = CK,$ |
| 5 | $\frac{d}{dt}[Pi] = r1 - r2,$ |
| 6 | $\frac{d}{dt}[AMPK] = -rc + JM_{13} + JM_{14} - JM_{15},$ |
| 7 | $\frac{d}{dt}[pAMPK] = rc + JM_{15} - JM_{13} - JM_{14},$ |
|  | Calcium and IP3 |
| 8 | $\frac{d}{dt}[Glut] = -200 * Glut,$ |
| 9 | $\frac{d}{dt}[Ca_{ER}] = (J_{SERCA} - J_{IP3R} - J_{RYR} - J_{ER,leak}) - J_{ER,Buff},$ |
| 10 | $\frac{d}{dt}[Ca_C] = -(J_{SERCA} - J_{IP3R} - J_{RYR} + J_{ER,leak}) + J_{PM} - J_{BuffCa} - J_{CaMBind},$ |
| 11 | $\frac{d}{dt}[w] = J_w,$ |
| 12 | $\frac{d}{dt}[Ri] = J_{Ri},$ |
| 13 | $\frac{d}{dt}[R2] = J_{R2},$ |
| 14 | $\frac{d}{dt}[DIM] = J_{DIM},$ |
| 15 | $\frac{d}{dt}[DAG] = J_{DAG},$ |
| 16 | $\frac{d}{dt}[DIMp] = J_{DIMp},$ |
| 17 | $\frac{d}{dt}[PKC] = J_{PKC},$ |
| 18 | $\frac{d}{dt}[IP3] = J_{IP} - k_{deg}(IP3 - IP3_0),$ |
| **Buffer and Calmodulin** |  |
| 19 | $\frac{d}{dt}[B] = -J_{BuffCa}$ |
| 20 | $\frac{d}{dt}[BCa] = +J_{BuffCa}$ |
| 21 | $\frac{d}{dt}[BuffER] = J_{ER,Buff}$ |
| 22 | $\frac{d}{dt}[CaM] = -J_{CaMBind} + J_{CaMDisc1}$ |
| 23 | $\frac{d}{dt}[CaCaM] = J_{CaMBind} - J_{CaMDisc}$ |
| 24 | $\frac{d}{dt}[CaMKK2] = -J_{CaMKK2,Act} + J_{CaMKK2,Deac}$ |
| 25 | $\frac{d}{dt}[CaMKK2_{act}] = J_{CaMKK2,Act} - J_{CaMKK2,Deac}$ |
| **Insulin System** |  |
| 26 | $\frac{d}{dt}pIR] = JM2 - JM1$ |
| 27 | $\frac{d}{dt}[pIR] = JM1 - JM2$ |
| 28 | $\frac{d}{dt}[IRS] = JM4 + JM16 - JM3 - JM15$ |
| 29 | $\frac{d}{dt}[pIRS] = JM3 - JM4$ |
| 30 | $\frac{d}{dt}[iIRS] = JM15 - JM16$ |
| 31 | $\frac{d}{dt}[AKT] = JM6 - JM5$ |
| 32 | $\frac{d}{dt}[pAKT] = JM5 - JM6$ |
| 33 | $\frac{d}{dt}[mTORC1] = JM8 - JM7 - JM13$ |
| 34 | $\frac{d}{dt}[pmTORC1] = JM7 - JM8$ |
| 35 | $\frac{d}{dt}[mTORC2] = JM10 - JM9 - JM14$ |
| 36 | $\frac{d}{dt}[pmTORC2] = JM9 - JM10$ |
| 37 | $\frac{d}{dt}[mTORC1 - DEPTOR] = JM13$ |
| 38 | $\frac{d}{dt}[mTORC2 - DEPTO] = JM14$ |
| 39 | $\frac{d}{dt}[DEPTOR] = JM12 - JM11 - JM13 - JM14$ |
| 40 | $\frac{d}{dt}[pDEPTOR] = JM11 - JM12$ |
| 41 | $\frac{d}{dt}[SIRT1] = JM19$ |
| 42 | $\frac{d}{dt}[ULK1] = JM21 - JM20$ |
| 43 | $\frac{d}{dt}[pULK1] = JM20 - JM21$ |
| **AMPA and NMDA** |  |
| 44 | $\frac{d}{dt}[NMDA_{C0}] = -JN1,$ |
| 45 | $\frac{d}{dt}[NMDA_{C1}] = +JN1 - JN2,$ |
| 46 | $\frac{d}{dt}[NMDA_{C2}] = +JN2 - JN3 - JN4,$ |
| 47 | $\frac{d}{dt}[NMDA_D] = JN3,$ |
| 48 | $\frac{d}{dt}[NMDA_O] = JN4,$ |
| 49 | $\frac{d}{dt}[AMPA_U] = -JA1,$ |
| 50 | $\frac{d}{dt}[AMPA_M] = +JA1 - JA2 - JA4,$ |
| 51 | $\frac{d}{dt}[AMPA_C] = JA2 - JA3 - JA5,$ |
| 52 | $\frac{d}{dt}[AMPA_O] = JA3 - JA6,$ |
| 53 | $\frac{d}{dt}[AMPA_{D1}] = JA4 - JA7,$ |
| 54 | $\frac{d}{dt}[AMPA_{D2}] = JA5 + JA7 - JA8,$ |
| 55 | $\frac{d}{dt}[AMPA_{D3}] = JA6 + JA8,$ |
| 56 | $[V] = -65 + BPAP + EPSP + AMPA_{EPSP},$ |

**Table 2:**
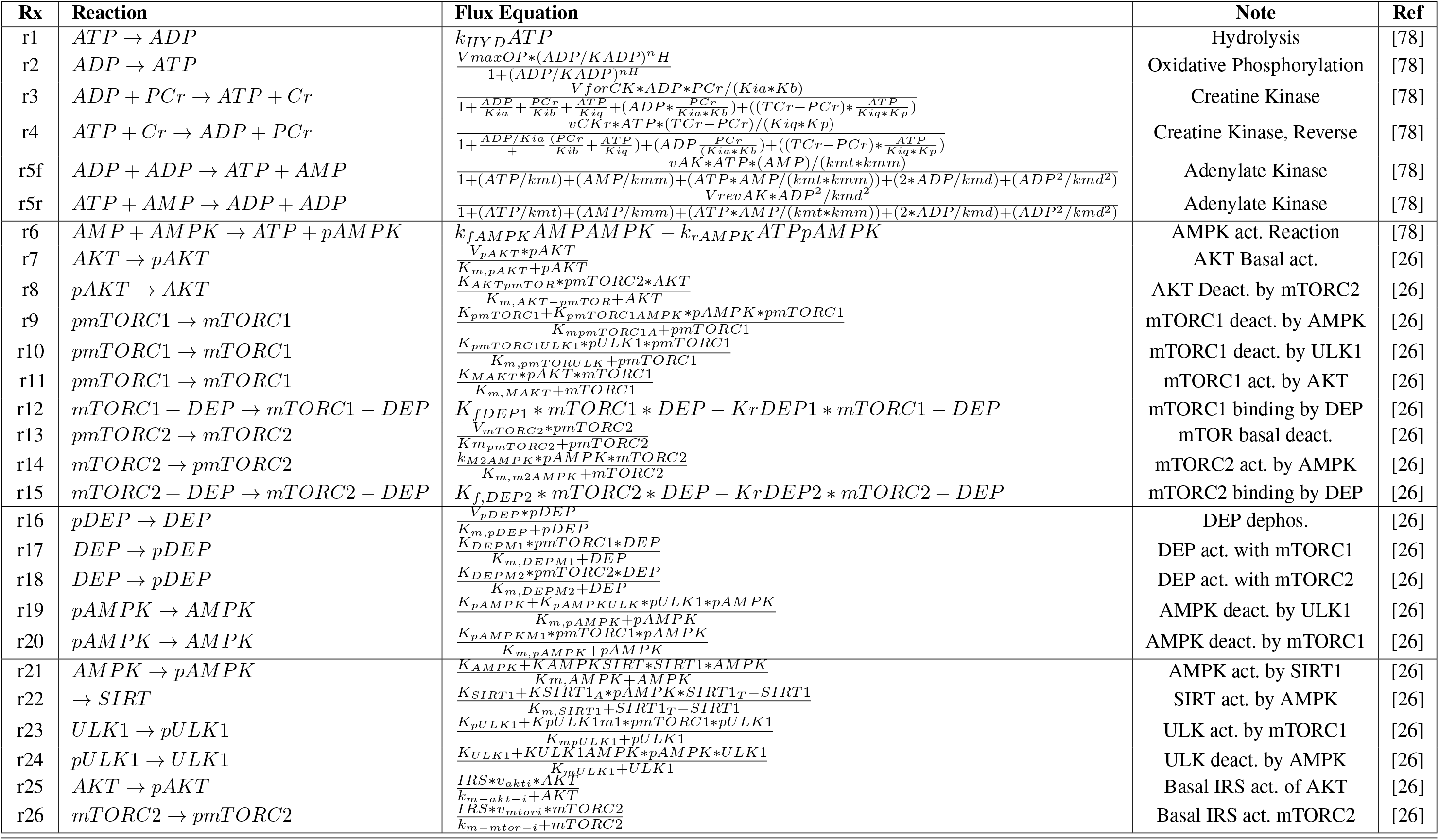
AMPK, mTOR, and Metabolism Reactions.

**Table 3:**
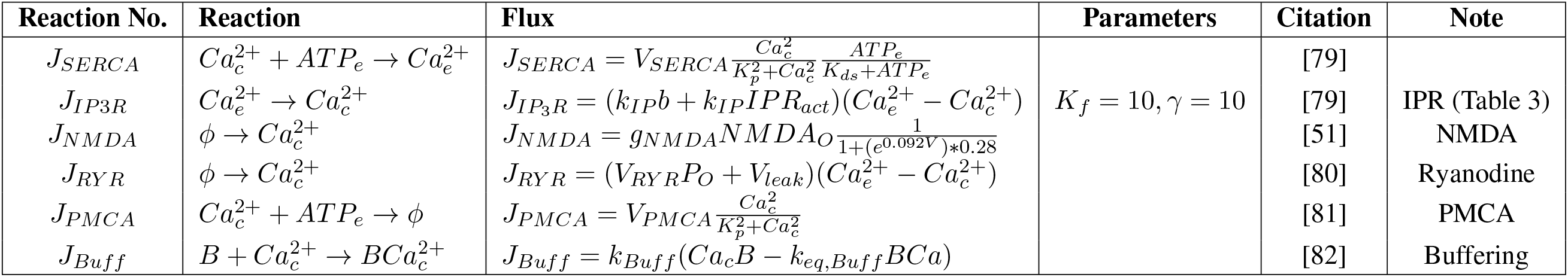
Calcium Submodel Reactions.

**Table 4:**
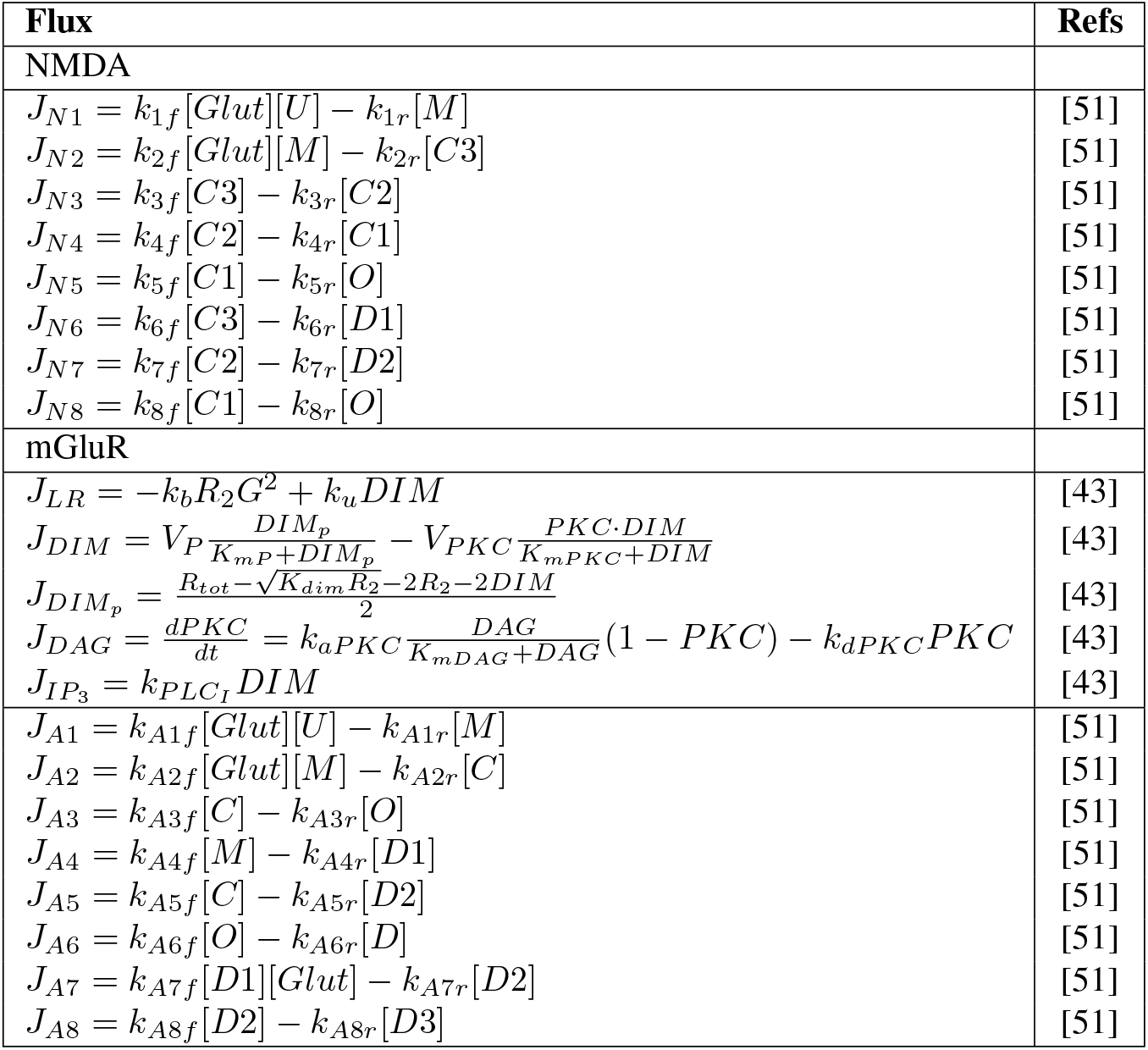
PSD Receptor Submodel Reactions.

**Table 5:** Parameter Table.

| # | Parameter | Value | Ref | # | Parameters | Value | Ref |
| --- | --- | --- | --- | --- | --- | --- | --- |
|  | Calcium Module |  |  |  |  |  |  |
| 31 | $k_1$ | 30 | [79] | 32 | $b$ | 0.001 | [79] [Altered] |
| 33 | $K_i$ | 5 $[\mu\text{M}]$ | [80] | 34 | $K_a$ | 3 $[\mu\text{M}]$ | [80] |
| 35 | $I_{Ra}$ | $EQN$ | [79] | 36 | $k_-$ | 0.02 $[1/\text{s}]$ | [79] |
| 37 | $k_-$ | 20 $[1/\mu\text{M}^4\text{s}]$ | [80] | 38 | $V_{DAG}$ | 0.0325 $[1/\mu\text{M}]$ | [80] |
| 39 | $k_b$ | 0.1 $[1/\mu\text{M}^2\text{s}]$ | [43] | 40 | $k_u$ | 2 $[1/\text{s}]$ | [43] |
| 41 | $V_p$ | 0.05 $[\mu\text{M}/\text{s}]$ | [43] | 42 | $K_{mp}$ | $5 \times 10^{-4}$ $[\mu\text{M}]$ | [43] |
| 43 | $V_{pkc}$ | 0.2 $[\mu\text{M}/\text{s}]$ | [43] | 44 | $K_{mpkc}$ | $5 \times 10^{-4}$ $[\mu\text{M}]$ | [43] |
| 45 | $k_{apkc}$ | 0.2 $[1/\text{s}]$ | [43] | 46 | $k_{dpkc}$ | 20 $[1/\text{s}]$ | [43] |
| 47 | $k_{plcI}$ | 3 $[1/\text{s}]$ | [43] | 48 | $k_{plcD}$ | 3 $[1/\text{s}]$ | [43] |
| 49 | $K_{mDAG}$ | 6 $[\mu\text{M}]$ | [43] | 50 | $V_{RYR}$ | 0.0050 $[1/\text{s}]$ | [80] |
| 51 | $K_{ar}$ | 0.0192 $[1/\text{s}]$ | [80] | 52 | $K_{br}$ | 0.2573 $[1/\text{s}]$ | [80] |
| 53 | $K_{cr}$ | 0.0571 $[1/\text{s}]$ | [80] | 54 | $K_{dr}$ | 0.1 $[1/\text{s}]$ | [80] |
| 55 | $V_{SERCA}$ | 120 $[\mu\text{M}/\text{s}]$ | [79] | 56 | $K_{pSERCA}$ | 6 $[\mu\text{M}]$ | [?] |
| 74 | $K_{m_{ext}}$ | 0.1 $[\mu\text{M}^2]$ | [81] | 75 | $k_{deg}$ | 5 $[1/\text{s}]$ | [79] |
| 76 | $k_{pmleak}$ | 0.5 $[1/\text{s}]$ | [81] | 77 | $k_{bcaf}$ | 0.1 $[1/(\mu\text{M s})]$ | [1] |
| 78 | $k_{bcar}$ | 1 $[1/\text{s}]$ | [1] | 79 | $k_{ERBf}$ | 50 $[1/(\mu\text{M s})]$ | [82] |
| 80 | $k_{ERBr}$ | 3 $[1/\text{s}]$ | [82] | 81 | $k_{erleak}$ | 0.1 $[1/\text{s}]$ | [To Sustain] |
| 99 | $k_{cam_f}$ | 0.001 $[1/(\mu\text{M s})]$ | [82] | 100 | $k_{cam_r}$ | 1 $[1/\text{s}]$ | [82] |
| 101 | $V_{m_{kk2}}$ | 0.01 $[\mu\text{M}/\text{s}]$ | [83] | 102 | $K_{m_{kk2}}$ | 0.01 $[\mu\text{M}]$ | [83] |
| 103 | $K_{a_{kk2}}$ | 0.005 $[\mu\text{M}]$ | [83] | 104 | $k_{f_{CaMDisc}}$ | 1 $[1/\text{s}]$ | [82] |

### Calcium dynamics

Intracellular calcium is a critical cellular second messenger that is closely linked to the induction of synaptic plasticity [42]. In this model, we source reactions established in legacy literature models of neuronal calcium [1–3,38,43]. Calcium influx is initiated by glutamate binding to receptors on the PSD described in receptor models. Calcium release from the endoplasmic reticulum (ER) is triggered through binding of IP3 to IP3R on the ER membrane, as well as calcium-induced calcium release by ryanodine receptors (RyR) [44]. These species are modeled as concentrations in different cellular compartments and described in the equations listed below:

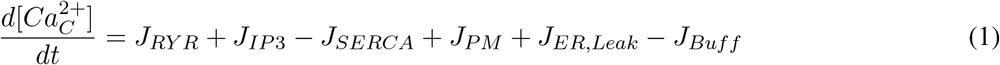

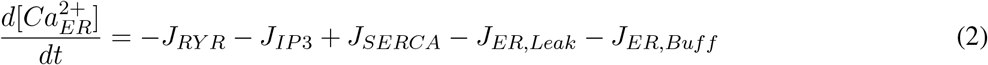

Here, 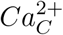 represents the calcium concentration in the cytosol and 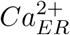 represents the ER calcium concentration. Flux components of these differential equations are represented by Ryanodine (*J_RYR_*), IP3 Receptor (*J*_*IP*3_), SERCA pump (*J_SERCA_*), plasma membrane calcium-atpase (*J_PM_*), ER calcium leak (*J_ER,leak_*), and buffering terms for the cytosol (*J_Buff_*) and ER (*J_ER,Buff_*) and are fully described in Table 3.

### ATP Production and Consumption

ATP is rapidly consumed during synaptic activation due to the export of ions as well as many housekeeping reactions involved in synaptic plasticity [13,14,19,45]. ATP is generated by glycolysis and oxidative phosphorylation. Glycolysis is a well-studied pathway in which cells metabolize glucose into pyruvate to generate ATP [16]. Computational models of glycolytic activity in neurons have provided predictions for neurodegeneration, pathology, and development [46–48]. Pyruvate, the end product of glycolysis, is oxidized in mitochondria; in our model, oxidative phosphorylation is a function of mitochondrial potential computed by fluxes of the electron transport chain, as described by Beard [49]. In addition, as oxidative phosphorylation is enhanced by high cellular calcium concentrations, we have incorporated this dependence with a Hill-type equation on the flux which represents mitochondrial energy production, *J_OP_*. In contrast, ATP is hydrolyzed to ADP to provide energy for many reactions in the cytosol. In our model, both ATP production and consumption are modeled as mass action kinetics with kinetic parameters from the literature [49]. However, the rates of energy consumption are dependent on synaptic signaling [12,13]. In this model, we assume that the active synaptic energy consumption is correlated with the activity of active transporters, for example, SERCA and PMCA. In addition to active energy consumption from synapses, global energy consumption through maintenance and housekeeping reactions add up to the overall energy consumption rate in the model. ATP, ADP and AMP are modeled in the cytosol as described in equations listed below:

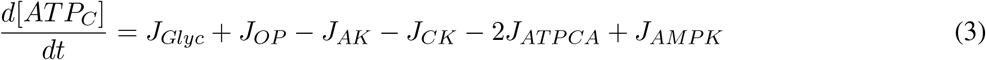

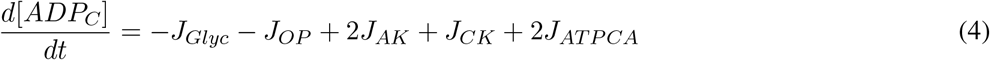

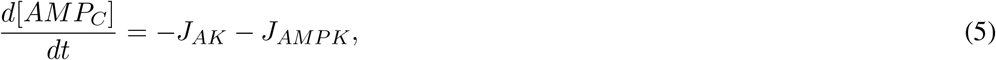

where ATP, ADP, and AMP are nucleotide concentrations over time. The fluxes modeled in these equations include: Glycolysis (*J_Glyc_*), Oxidative Phosphorylation (*J_OP_*), Adenylate Kinase (*J_AK_*), Creatine Kinase (*J_CK_*), ATP consumption by SERCA and PMCA pumps (*J_ATPCA_*), and AMPK activation (*J_AMPK_*) and are given in Table 2.

### Receptor models

Glutamate release into the synapse from the presynaptic cell is modeled as a series of step functions during the stimulus duration, with stimulus frequencies ranging from 0.1 to 100 Hz (Figure 4a-c). The decay of glutamate within the synapse is based upon time constants from experiments by Clements [50]. NMDAR and AMPAR cascades are modeled as a multi-state receptor model with a prescribed voltage [51]. These equations are included in Tables 2 to 4 and parameters in Tables 5 to 8.

**Table 6:** Parameter Table.

| # | Parameter | Value | Ref | # | Parameters | Value | Ref |
| --- | --- | --- | --- | --- | --- | --- | --- |
|  | PSD Module |  |  |  |  |  |  |
| 57 | $Rb$ | 5 $1/(\mu M s)$ | [51] | 58 | $Ru$ | 46.5 $[1/s]$ | [51] |
| 59 | $Rd$ | 8.4 $[1/s]$ | [51] | 60 | $Rr$ | 6.8 $[1/s]$ | [51] |
| 61 | $Ro$ | 46.5 $[1/s]$ | [51] | 62 | $Rc$ | 73.8 $[1/s]$ | [51] |
| 63 | $\tau_{ess}$ | 0.05 $[s]$ | [7] | 64 | $\tau_{esf}$ | 0.005 $[s]$ | [7] |
| 65 | $\tau_{bss}$ | 0.025 $[s]$ | [7] | 66 | $\tau_{bsf}$ | 0.003 $[s]$ | [7] |
| 67 | $\tau_{delay,bp}$ | 0.002 $[s]$ | [7] | 68 | $s$ | 10 $[mV]$ | [7] |
| 69 | $V_{rev}$ | -65 $[mV]$ | [3] | 70 | $N_{NMDA}$ | 1 | [51] |
| 71 | $BPAP_{max}$ | 40 $[mV]$ | [7] | 72 | $G_{NMDA}$ | $-65.6 \times 10^{-6} [nS]$ | [7] |
| 73 | $k_{ext}$ | $1.3 \times 10^{-6} [1/s]$ | [Altered] [81] | 82 | $G_{AMPA}$ | 15 $[nS]$ | [7] |
| 83 | $kAMPA_{1f}$ | 1.8 $[1/(\mu M s)]$ | [51] | 84 | $kAMPA_{2f}$ | 10 $[1/(\mu M s)]$ | [51] |
| 85 | $kAMPA_{3f}$ | $1.6 \times 10^{-4} [1/s]$ | [51] | 86 | $kAMPA_{4f}$ | $7 \times 10^2 [1/s]$ | [51] |
| 87 | $kAMPA_{5f}$ | $1 \times 10^2 [1/s]$ | [51] | 88 | $kAMPA_{6f}$ | $3 \times 10^2 [1/s]$ | [51] |
| 89 | $kAMPA_{7f}$ | 10 $[1/s]$ | [51] | 90 | $kAMPA_{8f}$ | $1.6 \times 10^4 [1/s]$ | [51] |
| 91 | $kAMPA_{1r}$ | $2.4 \times 10^3 [1/s]$ | [51] | 92 | $kAMPA_{2r}$ | $1 \times 10^4 [1/s]$ | [51] |
| 93 | $kAMPA_{3r}$ | $5 \times 10^3 [1/s]$ | [51] | 94 | $kAMPA_{4r}$ | $1.5 \times 10^2 [1/s]$ | [51] |
| 95 | $kAMPA_{5r}$ | 2.1 $[1/s]$ | [51] | 96 | $kAMPA_{6r}$ | 15 $[1/s]$ | [51] |
| 97 | $kAMPA_{7r}$ | $1 \times 10^3 [1/s]$ | [51] | 98 | $kAMPA_{8r}$ | $1.2 \times 10^4 [1/s]$ | [51] |

**Table 7:** Parameter Table.

| # | Parameter | Value | Ref | # | Parameters | Value | Ref |
| --- | --- | --- | --- | --- | --- | --- | --- |
|  | ATP Energetics |  |  |  |  |  |  |
| 1 | $K_{ADP}$ | 500 $[\mu M]$ | [78] | 2 | $VmaxOP$ | 0.5 $[mM/s]$ | [78] |
| 3 | $nH$ | 2.568 | [78] | 4 | $k_{HYD}$ | $2.6 \times 10^{-2} [mM/s]$ | [78] |
| 7 | $vAK$ | 14.66 $[mM/s]$ | [78] | 8 | $kmt$ | 0.27 $[mM]$ | [78] |
| 9 | $kmd$ | 0.35 $[mM]$ | [78] | 10 | $kmm$ | 0.32 $[mM]$ | [78] |
| 11 | $keqadk$ | 0.744 | [78] | 12 | $vmax20$ | $2 \times 10^{-2} [mM/s]$ | [78] |
| 13 | $vmax21$ | $1 \times 10^{-4}$ | [78] | 14 | $km20$ | 1.4 $[mM]$ | [78] |
| 15 | $km21$ | $6.7 \times 10^{-2} [mM]$ | [78] | 16 | $k12f$ | 1 $[1/(mMs)]$ | [78] |
| 17 | $k12r$ | $1.82 \times 10^{-2} [1/s]$ | [78] | 18 | $k13f$ | 1 $[1/(mMs)]$ | [78] |
| 19 | $k13r$ | $1.82 \times 10^{-2} [1/s]$ | [78] | 20 | $kaicar$ | $4 \times 10^{-4} [1/s]$ | [78] |
| 21 | $kAKT$ | $3 \times 10^{-3} [1/s]$ | [78] | 22 | $kcatAMPK$ | 34.56 $[1/s]$ | [78] |
| 23 | $V_{CK}$ | $1 \times 10^2$ | [78] | 24 | $Kb$ | 1.11 $[mM]$ | [78] |
| 25 | $Kia$ | 0.135 $[mM]$ | [78] | 26 | $Kib$ | 3.9 $[mM]$ | [78] |
| 27 | $Kiq$ | 3.5 $[mM]$ | [78] | 28 | $Kp(ATP)$ | 3.8 $[mM]$ | [78] |
| 29 | $KeqCK$ | $1.77 \times 10^2$ | [78] | 30 | $TCr$ | 39 $[mM]$ | [78] |

**Table 8:**
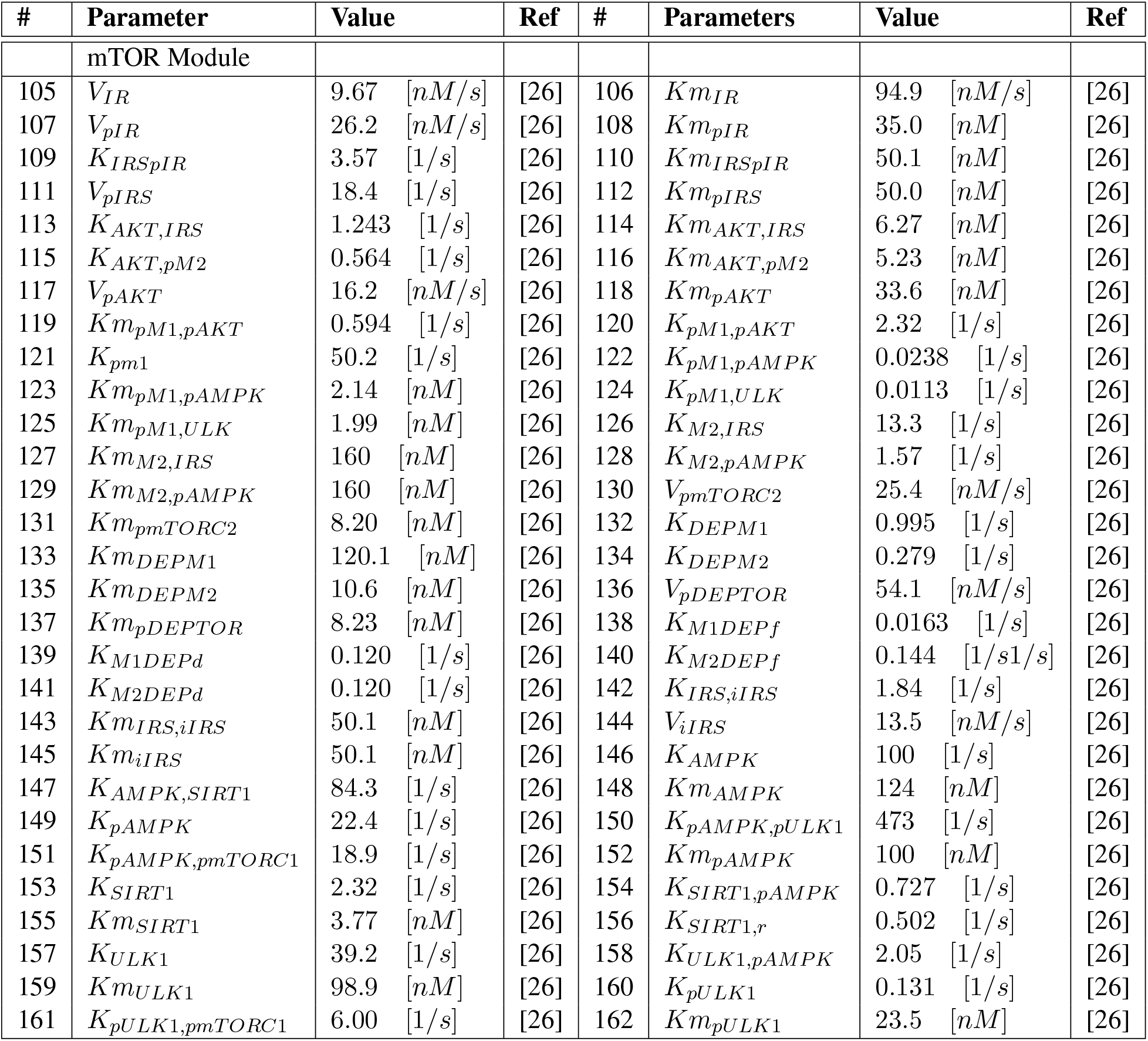
Parameter Table.

### AMPK and mTOR Activation

AMPK and mTOR are coupled in an intricate feedback loop involving several other protein kinases. This system is directly downstream of insulin receptor signaling and is derived from Sadria [26] (shown in Figure 1). Sadria et al. developed an AMPK and mTOR signaling pathway for activity in cancer cells, parameterized from experimental data of mTOR activation in adipocytes of type 2 diabetes patients [52]. We have adapted the signaling network and differential equations for species including mTORC1, mTORC2, AKT, ULK1, SIRT1, IRS, and have integrated metabolic activation of AMPK by AMP/ATP ratio.

### Mathematical Methods

The system of equations contains 60 species and 163 parameters. All equations were solved as a system of ordinary differential equations (ODE) using MATLAB’s built-in stiff solver, ode15s [53]. These equations were integrated with a maximum timestep of 0.1 s, relative integration tolerance of 10^-5^, using backward differentiation formulas with a maximum order of 4. Because each stimulus requires an instantaneous change in concentration and ODE solvers require a smooth, differentiable function, each stimulus was discretized to an individual ODE solution using the previous state as the subsequent initial condition, while glutamate concentration was increased to 0.1 mM. Before each pulse train simulation, the system was run to an approximate steady state, around 10,000 seconds with no glutamate stimulus. The results of this initial simulation are included in Supplemental Figure 1 and all model files are included in the public repository (https://github.com/aleung15/AMPKmTOR2022).

### Sensitivity Analysis

The dynamics of the model are dependent on both parameter values and initial conditions. The size and complexity of the system suggest nonlinear behavior such that changes in parameters for a single flux may lead to diverse response in the AMPK, AKT, mTORC1, and mTORC2 concentrations. To elucidate the role of each parameter in the system output we perform global sensitivity analysis. There are many methods of global sensitivity analysis, including correlation-based methods, variance-based methods, and derivative-based method [54]. In this work, we use a Latin Hypercube Sampling method to determine a sampling plan across a parameter range of 20% of the original parameter values [55]. Each parameter set is then simulated to an apparent steady-state and values for AMPK, AKT, mTORC1, and mTORC2 are compared to the default values. We then perform a partial correlation analysis to determine the Partial Rank Correlation Coefficient, which represents the linear dependence between the parameters and output variables. This approach is suitable for the analysis of the steady state simulation, since many of the non-linearities are introduced through the rapid glutamatergic and calcium signaling [56].

### Determination of system metrics

We use the following metrics to compare model outputs for different inputs: steady state, time to equilibrium, maximum amplitude, and area under the curve. Steady-state was determined by taking the derivative of phosphorylated AMPK, mTORC1, and mTORC2 with respect to time. After the mean magnitude of the derivative was computed to be lower than 1 × 10^-6^ μM/s for a period of 30 seconds, the species was determined to be at steady state. For the time to equilibrium, the same computation was done, however the time at which the derivative was lower than 1 × 10^-6^ μM/s was determined to be the time to equilibrium. Maximum amplitude was computed to be the percent change relative to the steady-state value for each condition. Finally, area under the curve (AUC) was numerically determined by the integrating the predicted phosphorylation time-course over a period of 200 seconds. In heatmaps Figures 7 to 9, the values of the AUC’s are taken relative to a system without any stimulus or changes in parameters, then normalized to the maximum AUC computed. This was done in order to reduce the range of values on the color bar to be more reflective of the changes in simulation conditions rather than the steady-state concentrations of the species.

## Results

Within a signaling synapse, the presynaptic neuron releases glutamate vesicles from low frequencies (0.1 to 1 Hz) to high frequencies (10 to 100 Hz). Low frequency signaling is often associated with long term depression and synaptic pruning, while high frequency signaling is believed to induce long term potentiation (LTP), the mechanism behind neuronal learning [57, 58]. Each pulse of glutamate triggers a calcium influx into the post synaptic site requiring a significant cellular energy cost in the form of ATP consumption for the restoration of resting ion potentials and various housekeeping processes (Figure 1b). As frequencies rise, there may be a critical point at which the ATP production from neuronal metabolism is overwhelmed by the energy demand. At this critical juncture, if the neuron is unable to adequately scale energy production, we hypothesize that the neuron may not be able to form sustain LTP and opt to undergo LTD. The metabolic plasticity, or the ability of the neuron to scale metabolic production to energetic demand, must be modeled alongside the closely coupled calcium signaling cascade to understand how the neuron is able to induce synaptic plasticity during periods of extreme energy stress. In what follows, we investigate the crosstalk between synaptic signaling and metabolic plasticity.

### Model constraints and parametric sensitivity analysis

We first analyzed the system behavior in the absence of any glutamate stimulus to investigate how the coupled signaling networks behave. After a brief initialization period, we see a rapid equilibration of all species to an apparent steady state, shown in Supplemental Figure 1. Many trajectories on the short, millisecond to second, timescale equilibrate nearly instantaneously including calcium, receptors, AMP/ATP, and AMPK. However, several species, like mTORC1, mTORC2, and IP3, take much longer to reach an equilibrium, on the second to minute timescale. Since the initial conditions are misaligned with the equilibrium states determined by key parameters, there is a brief initialization period before equilibration. For example, as the AMP/ATP ratio is much higher than the equilibrium value, this induces AMPK activity which leads to mTOR and AKT activity. Nevertheless, the resulting transient behavior fades after 200 seconds as the dampened oscillations reduce to a fixed point, representing the system at rest. All subsequent simulations use this rest state as initial conditions before stimulus.

The steady state behavior of the model is dependent on both parameters and initial conditions. However, due to the complexity of the pathway and the large amount of species and parameters, not all parameters have an equivalent effect on the end result of the model. To understand the complexity of the biological system posed in Figure 1 we first attempted to reproduce findings of experimental works. In Figure 2a, we compared the simulation of AMPK, mTORC1, and AKT phosphorylation relative to steady state to the experimental results of [23]. Our model predictions under 10 Hz of glutamate stimulus align well with their experimental values of differentiated primary neurons stimulated with Bic/4-AP protocol. However, to characterize the uncertainty of the model predictions, we then quantified the effect of parameters on model output.

**Figure 2:**
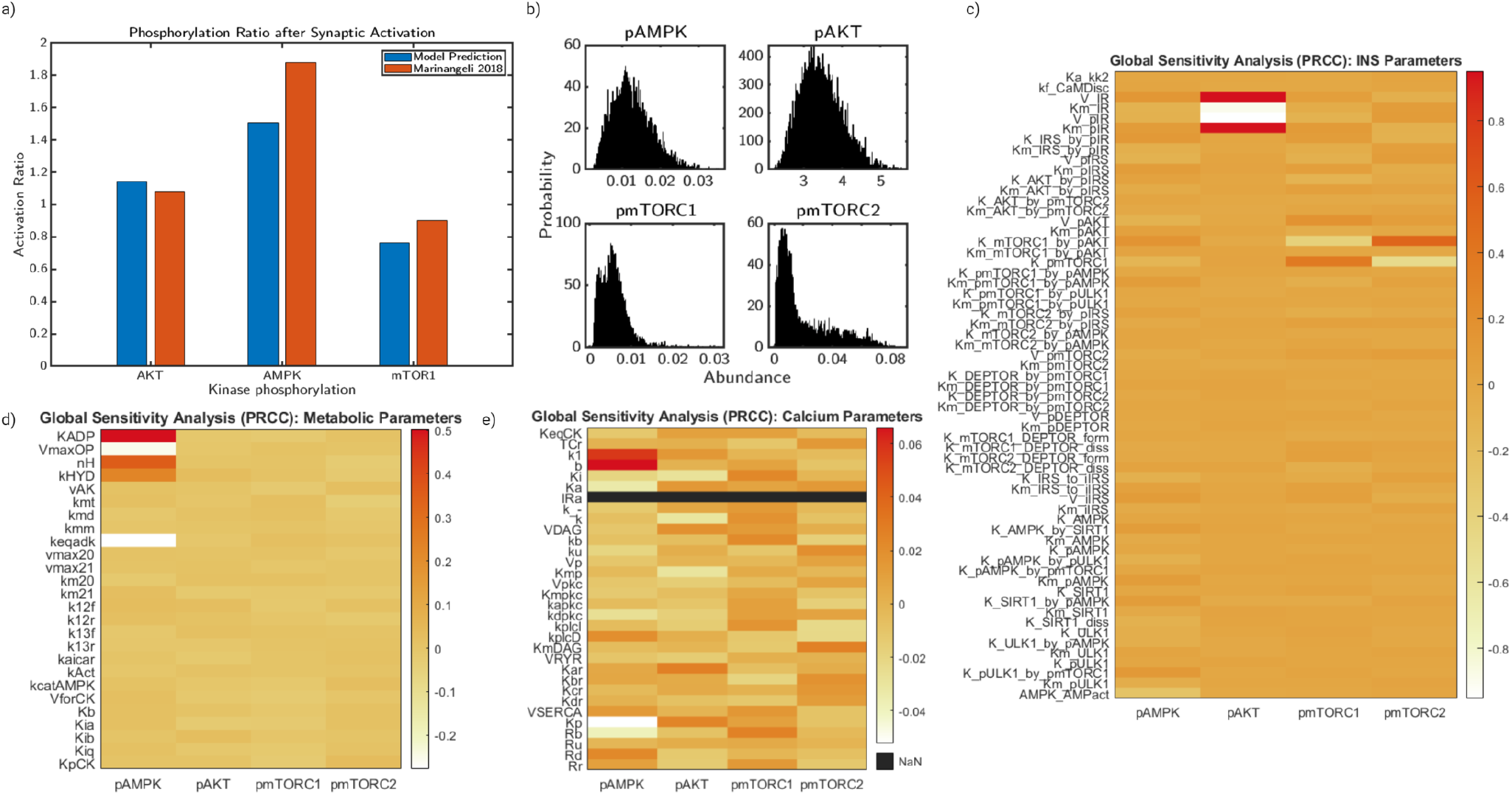
Sensitivity analysis reveal key parameters regulating system behavior. The model describing AMPK and mTOR phosphorylation due to glutamate-stimulated calcium influx contains 163 parameters and 60 equations in a well-mixed model, **a)** Comparisons between model predictions of phos-phorylation ratios relative to initial state after 10 Hz synaptic activation and experimental results from Marinangeli et al. 2018 primary neuron cells stimulated via Biciculin/4AP protocol, **b)** Probability density functions of steady-state concentrations of pAMPK, pAKT, pmTORC1, and pmTORC2 resulting from a global sensitivity analysis of 10,000 parameter values for each 163 parameters in a 20% range, c) Heatmap of PRCC values describing the correlation between parameter value and system output for AMPK, AKT, mTORC1, and mTORC2 phosphorylation for a subset of parameters belonging to the insulin signaling system, **d)** Heatmap of PRCC values describing the correlation between parameter value and system output for AMPK, AKT, mTORC1, and mTORC2 phosphorylation for a subset of parameters belonging to the neuronal metabolism system, **e)** Heatmap of PRCC values describing the correlation between parameter value and system output for AMPK, AKT, mTORC1, and mTORC2 phosphorylation for a subset of parameters belonging to the calcium signaling system.

Through global sensitivity analysis of the system using the PRCC method, we obtained correlation values of each parameter to the model predictions of AMPK, AKT, mTORC1, and mTORC2 phosphorylation. In Figure 2b, we show histograms of the steady-state values of the model predictions as a probability density function in which the x-axis denotes the steady state concentration and the y-axis denotes the number of parameter sets that predict the steady state. For AMPK concentration, the predictions are well distributed with a mean value of 0.02 μM. AKT also showed similar characteristics, with a mean value of 4.5 nM. This is consistent with expected model behavior as AKT is directly downstream of AMPK. mTORC1 and mTORC2 are both skewed heavily left with mean values of 0.012 and 0.07 μM, respectively. In Figure 2c-e, we plot heatmaps for the PRCC values of each parameter in the model and the effect on steady-state values of AMPK, AKT, mTORC1, and mTORC2 phosphorylation. We grouped the parameters into sets corresponding to their most relevant biological pathway. Figure 2c corresponds to the parameters most closely related to insulin receptor signaling, originally derived in [26]. While most values are close to the mean value, parameters downstream of AKT appear to have the most influence on mTORC1 and mTORC2. AKT is most strongly impacted by V_*IR*_, a parameter which represents the intercellular activity of the insulin receptor signaling and will be investigated.

Next, in Figure 2d, we plot the PRCC heatmap for the parameters associated with metabolism. For this segment of parameters, AMPK is most strongly impacted by oxidative phosphorylation parameters and the AMPK activation parameter. AKT, mTORC1, and mTORC2 are not strongly influenced by metabolic parameters at steady-state. Finally, in Figure 2e, we plot the PRCC heatmap for calcium related parameter values. The magnitude of these terms are very small relative to the effect shown in Figure 2c-d. Overall, we found that the model is very sensitive to parameters corresponding to oxidative phosphorylation, ATP consumption, and insulin signaling. While this implies that the model output is not particularly sensitive to calcium values at steady-state, we now aim to show how the system behaves under glutamatergic stimulus.

### Low frequency neuronal stimulus reveals low amplitude transient behavior

After establishing the steady state behavior in the absence of stimulus (Supplemental Figure 1), we perturb the system with a 1 Hz glutamate pulse for 50 seconds, which is representative of typical LTD induction protocol [59]. Glutamate stimulus induces a calcium influx (Figure 3a) into the cytosol through the glutamatergic receptors, which consequently induces ATP consumption related to ion transport, increases the AMP/ATP ratio (Figure 3b), and therefore AMPK activation. This AMPK activation (Figure 3c) then leads to an initial increase of mTORC1 (Figure 3d), mTORC2 (Figure 3e), and ULK1 (Figure 3f).

**Figure 3:**
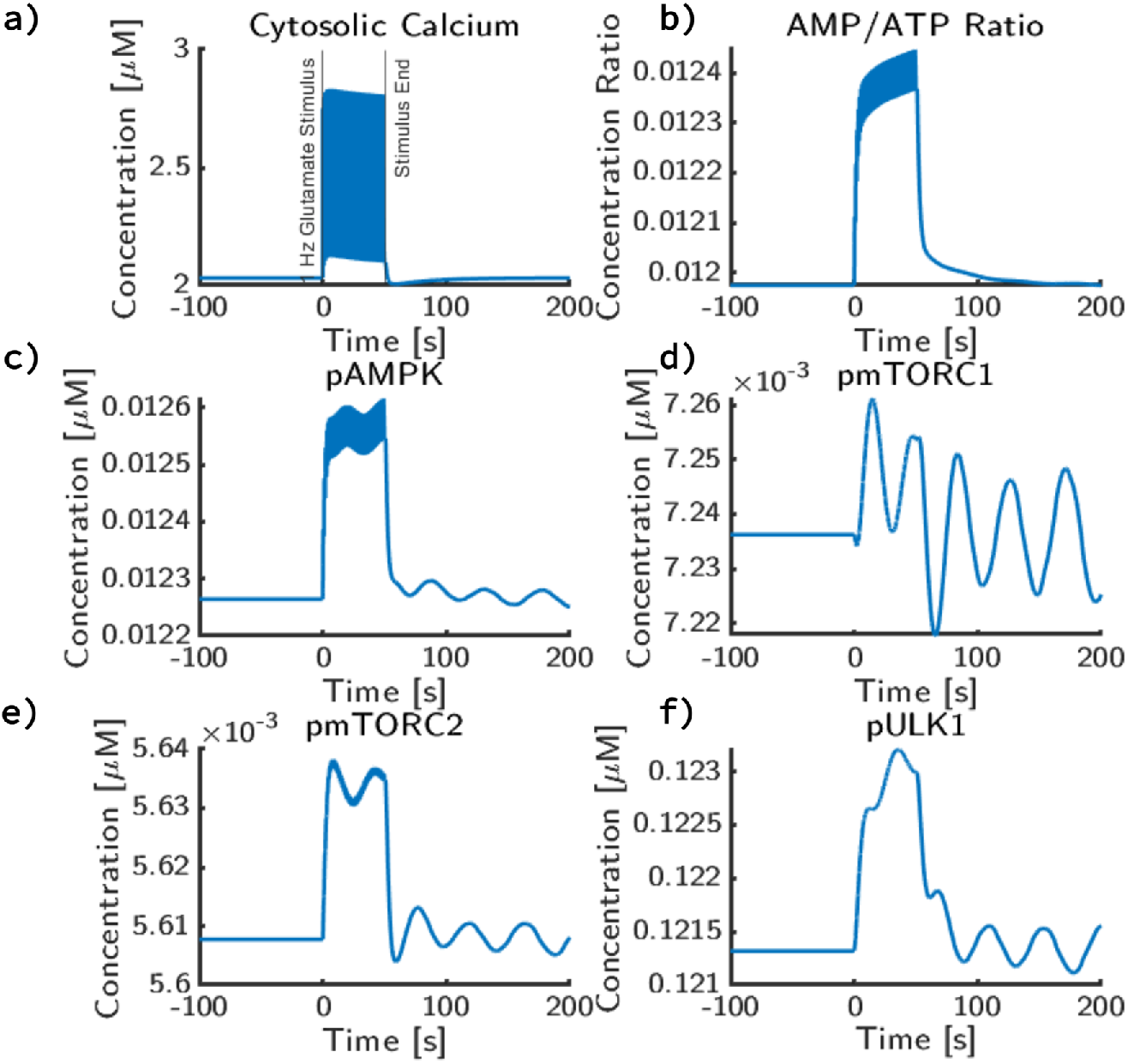
Effect of a simple 1 Hz glutamate on AMPK/mTORC dynamics. At t=0s, a 1 Hz pulse train over 50 seconds is applied to the system. Before stimulus, the system was allowed to reach a steady state. During each pulse, 100 μM of glutamate is applied, which decays with a rate constant of 200 ms. After the pulse train, the system was allowed to return to an apparent equilibrium with no additional glutamate input. Concentration trajectories for **a)** cytosolic calcium concentration, **b)** cytosolic AMP/ATP ratio, **c)** cytosolic phosphorylated AMPK concentration, **d)** phosphorylated mTORC1 concentration, **e)** phosphorylated mTORC2 concentration, **f)** phosphorylated ULK1 concentration are shown.

**Figure 4:**
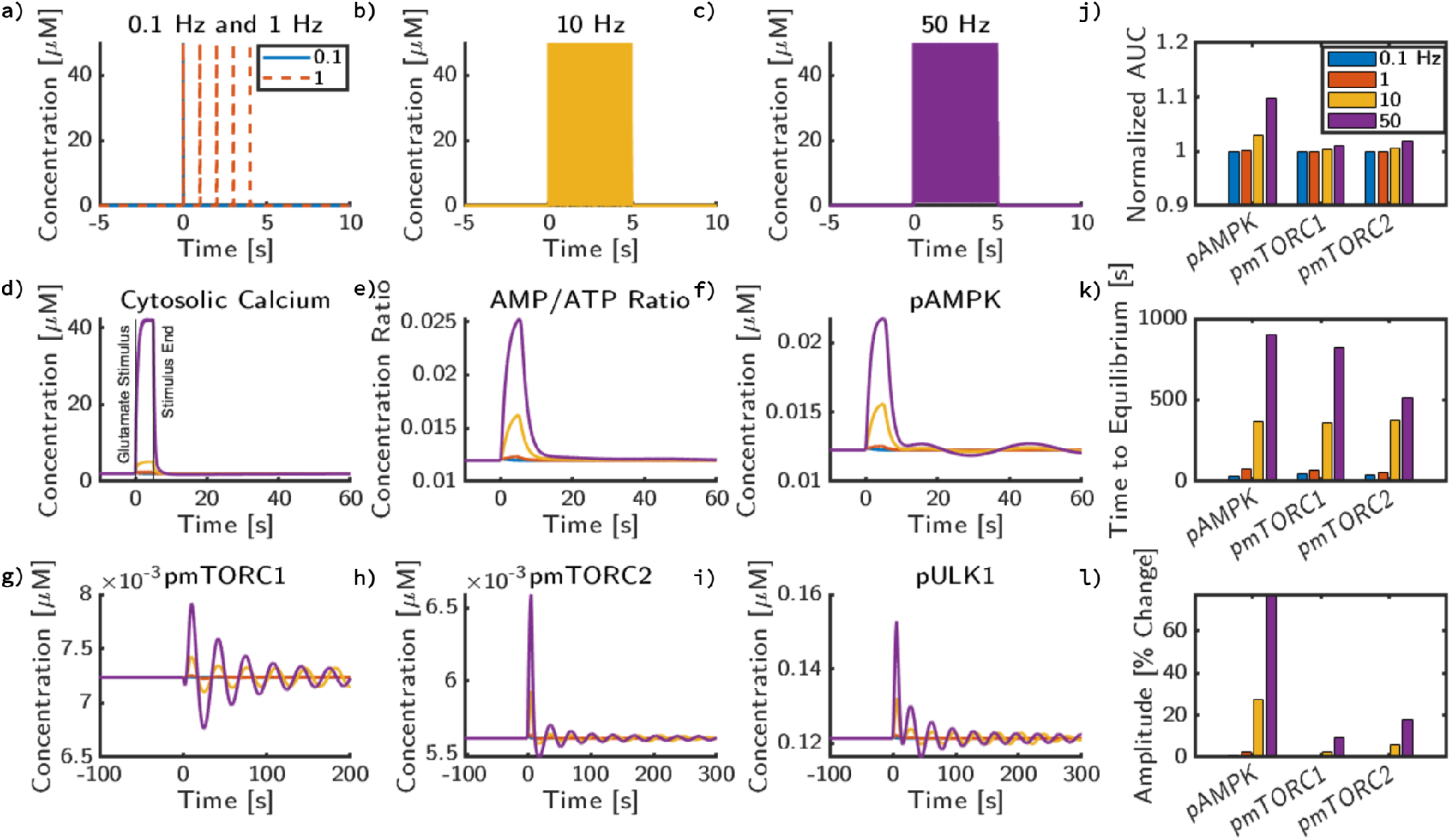
mTOR oscillations are stimulus frequency dependent. The system was stimulated with four different glutamate frequencies: **a)** 0.1 Hz (Blue), 1 Hz (Orange), **b)** 10 Hz (Yellow), **c)** 50 Hz (Purple). After stimulus, simulations return to an apparent equilibrium with no additional glutamate input. Trajectories are shown for **d)** cytosolic calcium concentration, **e)** cytosolic AMP/ATP ratio, **f)** phosphorylated AMPK concentration, **g)** phosphorylated mTORC1 concentration, **h)** phosphorylated mTORC2 concentration, **i)** phosphorylated ULK1 concentration. Quantitative metrics for AMPK, mTORC1, and mTORC2 in response to pulse trains of glutamate stimulus are also shown: **j)** Area under the curve (AUC) relative to the system without stimulus applied over the same integration window (100 seconds), **k)** time to reach equilibrium, and **l)** amplitude change quantified as percent change from steady state value.

Cytosolic calcium attains a peak calcium concentration of 4 μM and decays rapidly to the baseline concentration of 2 μM (Figure 3a). Due to the increased energy consumption associated with the calcium pumps, AMP/ATP ratio increases during synaptic signaling to a peak value of 0.0124 μM from its baseline value of 0.012 μM (Figure 3b). We note that both calcium and the AMP/ATP ratio (Figure 3b) match the frequency of the input glutamate. pAMPK also shows a corresponding dynamic (Figure 3c); phosphorylated AMPK increases by 19% but returns to baseline activation. However, instead of solely being influenced by glutamatergic signaling, AMPK also recieves signals from the insulin system, thus has longer timescale oscillations. The magnitude of change for critical terms like mTORC1, mTORC2, and ULK1 are fairly small during 1 Hz stimulus of glutamate. In response to this AMPK activation, mTORC1 and mTORC2 both deviate a maximum of of 0.5% but the trajectories observed are different; mTORC2 during stimulus reaches a higher peak value relative to its baseline oscillations while stimulus appears to lead mTORC1 to only slightly increase its amplitude (Figure 3d,e). Changes in ULK1 follow similar trajectories to mTORC2 with a 1.2% change in activation decaying shortly after stimulus ends back to unstimulated values (Figure 3f).

While the oscillation magnitudes are quite low in this case, it must be noted that the stimulus profile is most associated with LTD and therefore the energy demand and subsequent activation of downstream kinases of AMPK is expected to be low. However, this result is notable because it shows the cross-talk between two disparate signaling cascades through AMPK as a common vector. Through calcium and energy consumption, neuronal stimulus leads to direct changes in AMPK activation that influence the activity of mTORC1 and mTORC2. In the case of low frequency stimulus and base parameter values, there is little influence, but these results suggest that signaling frequency may impact this crosstalk.

### Effect of glutamate stimulus frequency on AMPK/mTOR pathway

We next investigated the effect of glutamate stimulus frequency on the activation profiles of key model outputs. The glutamate input stimuli, shown in Figure 4a-c, was varied from 0.1 Hz to 50 Hz, reflective of the range of potential stimulus frequencies in a signaling neuron [58]. While in Figure 3, we show only minor changes in state and overall signaling effect, as stimulus frequency increases, cytosolic calcium concentration and ATP consumption also increase (Figure 4d-e). The change in calcium concentration is dependent on the frequency of glutamate stimulus (Figure 4d); as stimulus frequency increases, calcium amplitude increases and the higher frequencies are filtered out (also see [2, 60]). There is a corresponding ATP consumption that leads to an increase in the AMP/ATP ratio (Figure 4e), which leads to AMPK activation (Figure 4f). In low levels of stimulus frequency, mTORC1 (Figure 4g) and mTORC2 (Figure 4h) oscillate around the baseline value. When AMPK is activated in higher magnitudes, we observe a large initial perturbation from the baseline oscillatory pattern that results in damped oscillations that eventually return to equilibrium concentrations. The maximum amplitude of the deviations from equilibrium of mTORC1, mTORC2, and ULK1 (Figure 4g-i) are dependent on the amplitude of AMPK activation, and therefore glutamate stimulus frequency. Increasing stimulus frequency does not appear to significantly lead to a phase shift in frequency for any species, but most notably increases the initial magnitude of oscillations.

The AUC over 100 seconds (Figure 4j) represents the total amount of a species within the system during short-term response to signaling pulse trains. Since the concentrations of each species can be orders of magnitude apart, we normalized each AUC to the baseline activity profiles to obtain a relative AUC value. In Figure 4j, we observed a frequency-dependent increase in the AUC of AMPK. However, for mTORC1 and mTORC2, only very minor increases in AUC were observed, 1 and 2%, respectively. At low frequency conditions (0.1 and 1 Hz), the AUC of all species does not change significantly. As frequency increases, AMPK activation increases proportionally to the highest stimulus frequency, where AMPK’s AUC increases by 10% after exposure to 50 Hz stimulus for 5 seconds. While the overall magnitude of change for mTORC1’s AUC is lower in comparison to pAMPK and mTORC2 for all frequencies, there is a slight increase as frequency increases.

Next, in Figure 4k and Figure 4l, we characterized the damped oscillatory behavior of the trajectories through its time to equilibrium and amplitude. All three species have a similar trend in which higher frequencies have longer times to equilibrium (TTE) and higher amplitudes. Another point of note is that the increase in TTE with respect to frequency is consistent up to 10 Hz, but 50 Hz shows a significant disparity between AMPK, mTORC1 and mTORC2. mTORC2 returns to equilibrium around 300 seconds faster than mTORC1 and 400 seconds faster than AMPK, which returns to equilibrium at 900 seconds. As for amplitudes (Figure 4l), AMPK maximum amplitude is significantly higher due to direct activation from AMP/ATP ratio. mTORC1 has the lowest increase in amplitude due to high frequency stimulus. The maximum amplitude for mTORC2 is higher than mTORC1, but even at 50 Hz, only 17.5 % increase over steady state. Overall, there is a separation between high frequency and low frequency stimulus in the observed trajectories and metrics. While the predicted AUC, TTE, and amplitude are similar between 0.1 and 1 Hz stimulus, the model predicts up to a 10 % increase in AUC for high frequency stimulus and a significant increase in observed oscillation duration.

### Increasing basal energy consumption increases AMPK activation

While calcium and other ion transport are known to be primary drivers of energy consumption in neurons, there are additional energy consuming processes that require ATP in the dendritic spine. For example, it is thought that actin, which is abundant in dendritic spines, is one of the main non-signaling ATP sinks through actin polymerization and remodeling [61]. Additionally, protein and lipid turnover in neurons can consume up to 25% of the total ATP consumed in brain tissue [14,62].

To explore the interaction between the basal energy consumption rate and the glutamate stimulus on AMPK signaling, we designed a series of simulations using our model. In our model, energy consumption is simplified as a lumped parameter representing ATP hydrolysis in the cytosol, *k_hyd_*. We varied this parameter to capture the effect of different energy consumption rates. We held our glutamate stimulus to 10 Hz because this stimulus approximates the threshold of frequency necessary to trigger change in mTOR (Figure 4). In this set of simulations, all variations had the same set of initial conditions, but at t=0, *k_hyd_* was changed in addition to glutamate pulse trains of 10 Hz.

We found that varying *k_hyd_* had led to direct changes in the peak and steady state behavior of AMP/ATP ratio (Figure 5a). There was no change in calcium dynamics since the calcium influx depends on the NMDAR fluxes, which are not dependent on hydrolysis rates. Compared to the baseline value, a 10-fold decrease in *k_hyd_* decreases the AMP/ATP ratio by a small amount but a 3-fold increase in k_hyd_ increases AMP/ATP ratio dramatically. This behavior is reflected in pAMPK (Figure 5b), where a nonlinear increase in pAMPK levels is seen for a 3-fold increase in the energy consumption rate. Interestingly, changing *k_hyd_* actually changes the oscillation pattern for pmTORC1 (Figure 5c). Increase in the energy consumption rate translates the damped oscillations observed in pmTORC1 at lower energy consumption rates into a stable steady state, without any oscillations, at high energy consumption rates. This effect is also seen in the dynamics of pmTORC2 (Figure 5d) and pULK1 (Figure 5e) except that increasing basal energy consumption increases the level of pmTORC2 and pULK1. Further analysis of these variations reveals that the steady state depends on the basal energy consumption (Figure 5f).

**Figure 5:**
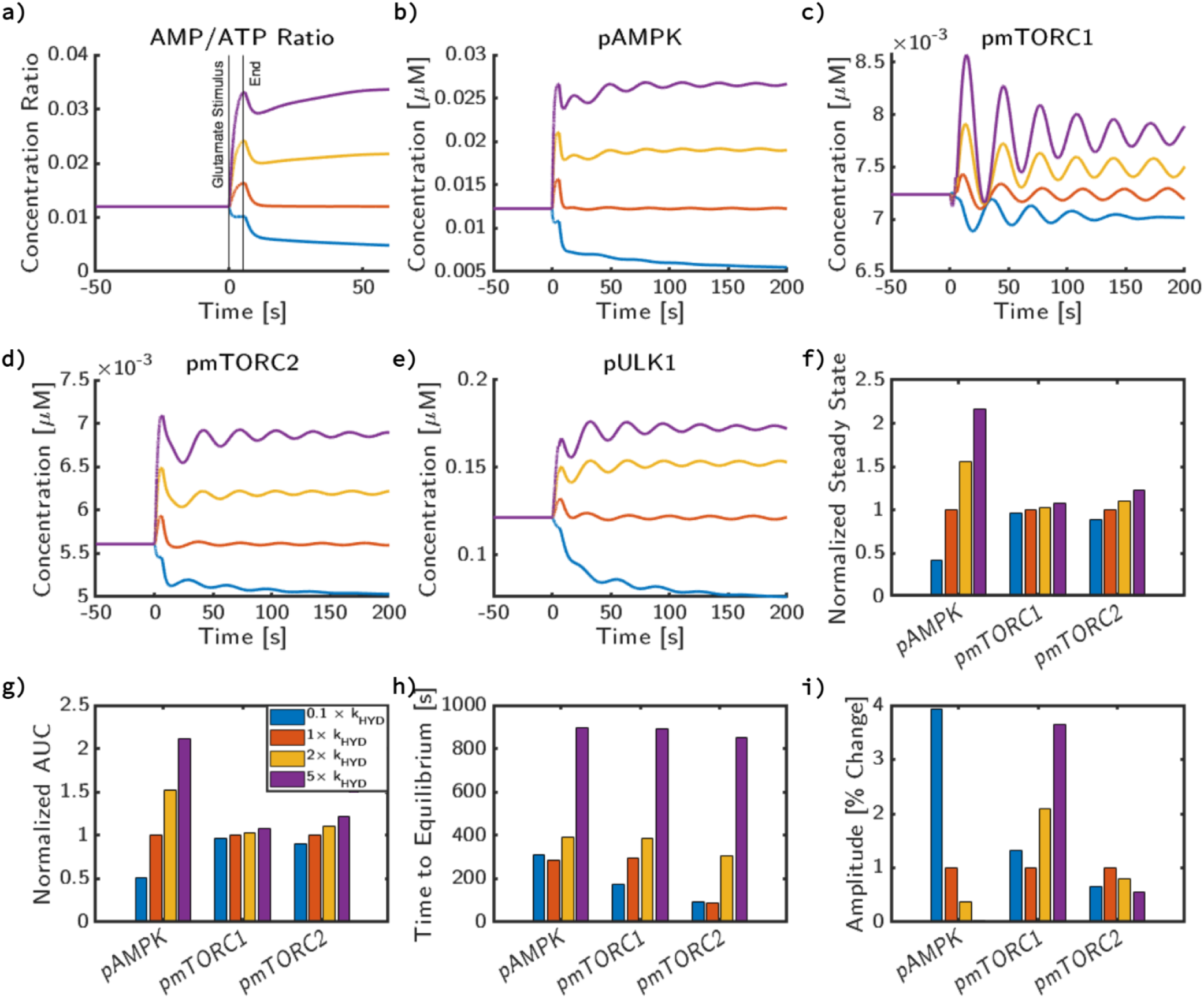
Cellular metabolic rate influences steady state behavior of AMPK and mTOR independent of calcium. We apply a 10 Hz glutamate stimulus for 5 seconds, however at t=0s, we also change the value of baseline energy consumption throughout the cell and plot concentration trajectories for **a)** AMP/ATP, **b)** active, phosphorylated AMPK, **c)** active, phosphorylated mTORC1, **d)** active, phosphorylated mTORC2, **e)** active, phosphorylated pULK1. Additionally, in **f)**, we compare the changes in steady state for pAMPK,mTORC1, and mTORC2 with respect to changes in hydrolysis rate. In **g)**, we then compare how this change impacts AUC, normalized to the AUC of simulation with the base value of the parameter. In **h)**, we plot the time to reach equilibrium and in **i)**, the relative magnitude of the first peak of AMPK, mTORC1, and mTORC2 to its new steady state value as a result of energy consumption from glutamatergic stimulus.

Next, we characterize the effect of hydrolysis rates on the dynamic features of AMPK, mTORC1, and mTORC2 signaling by plotting the normalized AUC, TTE, and percent amplitude (Figure 5g-i). For AMPK phosphorylation, there is an increase in AUC with increasing *k_hyd_*. From the baseline value of *k_hyd_*, reducing *k_hyd_* to a tenth of its original value roughly produces an AUC of half. However, the increase of *k_hyd_* to 2 × *k_hyd_* and 3 × *k_hyd_* increases the relative AUC by 50% and 120%, respectively. For mTORC1, while the increase in AUC with increasing *k_hyd_* is apparent, the magnitude of increase is not as high when compared to AMPK and is within a few percent of the steady state value. This trend can qualitatively be observed in Figure 5c, as the variations of *k_hyd_* from 0.1 to 2 × *k_hyd_* have roughly the same trajectories, but the variation to 3 × *k_hyd_* increases the steady state substantially. For mTORC2, the AUC increase from 0.1 to 1 × *k_hyd_* is minor relative to the change in *k_hyd_*, however 3 × *k_hyd_* produces a 20% jump in AUC. 2 × *k_hyd_* and 3 × *k_hyd_* both have proportional responses relative to the change in *k_hyd_*.

The time to equilibrium of AMPK increases slightly with both increasing and decreasing *k_hyd_* from baseline value. This is surprising, because for most metrics there appears to be a consistent trend between hydrolysis rate and metric value. The mTORC1 and mTORC2 TTE both follow similar trends of monotonic increase relative to *k_hyd_*. This could be caused by the 0.1 × *k_hyd_* equilibrium (blue curve) being further away from the initial state than the 1 × *k_hyd_* case (represented by the orange curve).

Finally, the amplitude change depends on the magnitude of *k_hyd_* since, as shown in Figure 5f, the steady state is dependent on *k_hyd_*. Therefore, for AMPK, as *k_hyd_* increases, the percent change in amplitude decreases. For mTORC1, there is a competing effect – as *k_hyd_* increases, the magnitude of deviation change increases, but the equilibrium values decrease. For mTORC2, as the steady state values increase with increasing *k_hyd_* but the percent change of amplitude decreases. Thus, we found that the stability features of the AMPK/mTOR pathway depend on the basal rate of ATP hydrolysis – increasing ATP consumption allows the system to transition from an oscillatory state to a stable steady state.

### Insulin signaling and metabolic signaling interact to govern stability features

Thus far, we have explored the impact of glutamate stimulus and energy consumption on AMPK and mTOR activation. We next investigated the role of the insulin response pathway and its crosstalk with the glutamate pathway. Insulin receptor signaling is a primary input in metabolism and influences the phosphorylation state of AMPK, mTORC1, and mTORC2. In the model originally developed by Sadria [26], the insulin receptor substrate (IRS) directly interacts with AMPK and mTORC2, resulting in regimes of oscillations dependent on the rate of IRS phosphorylation. Since we included this pathway in our model, we now explore the feedback between the IRS pathway and AMPK pathway through the energy consumption rate. In our model, the parameters controlling the IRS phosphorylation rate and energy consumption rate are, V_*IR*_ and *k_hyd_*, respectively. In Figure 2c-e, parameters corresponding to V_*IR*_ and *k_hyd_* had significant control over the steady state concentrations of AMPK, mTORC1, and mTORC2 under no glutamate stimulus. Here, we investigated how V_*IR*_, *k_hyd_*, in conjunction with glutamate stimulus impact the AMPK-mTOR pathway.

In Figure 6, we show how the stability of the model is dependent on the magnitude of *k_hyd_* and V_*IR*_. First, for hydrolysis (Figure 6a,b), we select two values of V_*IR*_ to hold constant (5.7368 mM/s yellow and 0.01 mM/s in purple) and vary the rates of *k_hyd_* (1000 values between 1 × 10^-4^ to 1 mM/s). We simulate the system in the absence stimulus until equilibrium for each parameter combination and compute the minima and maxima as a function of V_*IR*_. We observed that for the lower value of V_*IR*_, increasing *k_hyd_* gives a single unique steady state for both mTORC1 and mTORC2 (Figure 6a, b). For higher values of V_*IR*_, we found that the stability behavior has an oscillatory regime for low values of *k_hyd_* and a single steady state for high values of *k_hyd_*. Thus, the hydrolysis rate alone can alter the stability features of the coupled system. Next, to obtain the relation between stability and V_*IR*_ values, we select two values of *k_hyd_* and vary with 60 values of V_*IR*_ (from 0.1 to 20 mM/s), shown in Figure 6c,d. In Figure 6c, we compute the minima and maxima as a function of V_*IR*_ to show the stability features for two values of *k_hyd_*, 0.0001 (Blue) and 0.149 (Red) [mM/s]. For low and high values of V_*IR*_, minima and maxima curves converge to a singular value. However, there is a range of V_*IR*_ values in which the curves diverge. In this range, we see oscillations similar to those seen in the equilibrium states of Figure 3c-f. Values of V_*IR*_ that result in oscillatory behavior are correlated with hydrolysis; as *k_hyd_* increases, a smaller range of V_*IR*_ produces oscillations. Compared to a low value of *k_hyd_* (blue, 0.0001 [mM/s]), a high value of *k_hyd_* (red, 0.149 [mM/s]) shows a significantly smaller range of values, as well as a much smaller range of oscillations. This same approach is taken with Figure 6d for mTORC2.

**Figure 6:**
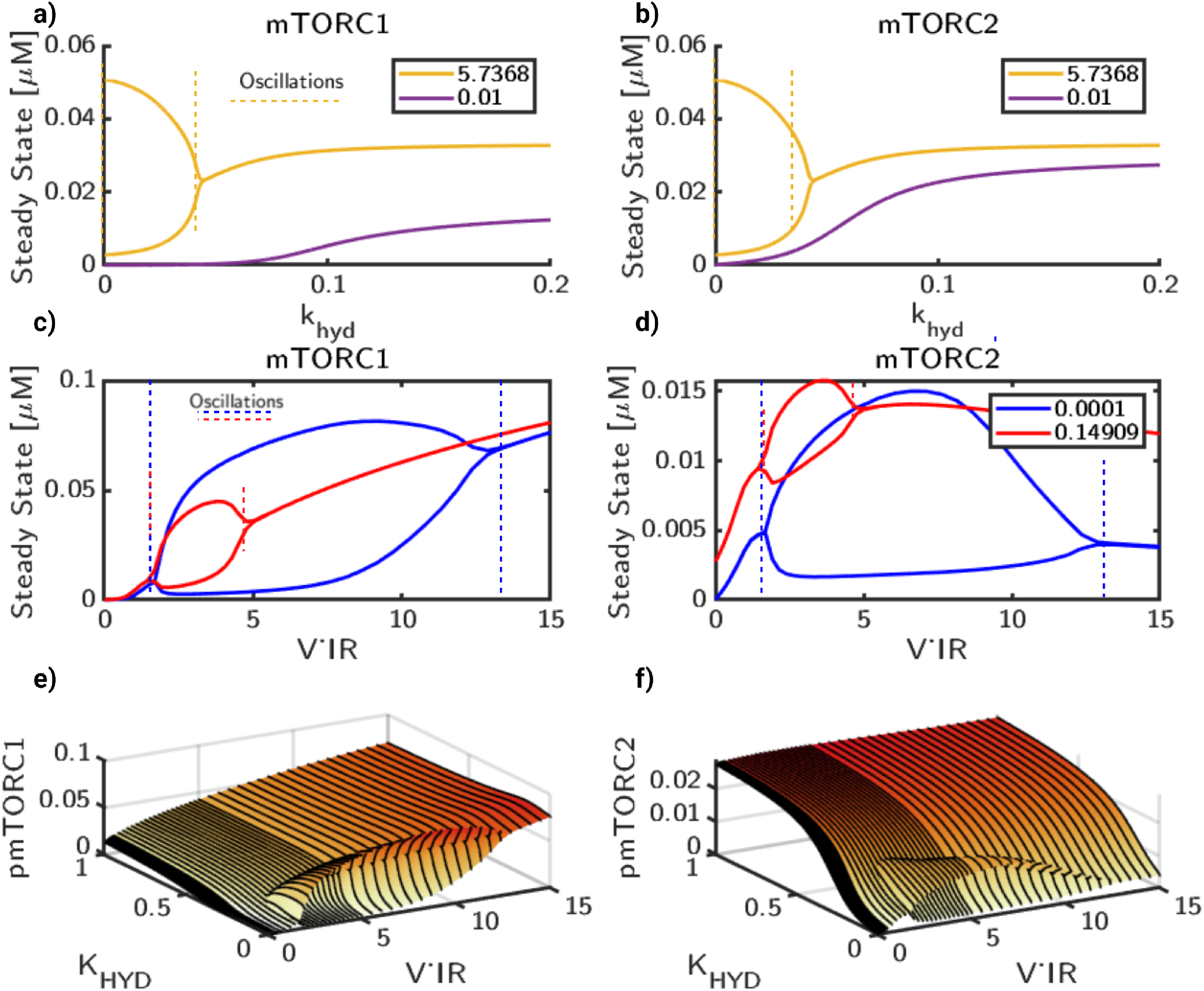
Oscillatory regimes for mTOR activation are dependent on both metabolic activity and IRS. System displays oscillatory behavior dependent on both *k_hyd_* and V_*IR*_ rates. We show the dependence of **a)** mTORC1 and **b)** mTORC2 stability on *k_hyd_* by selecting two values of V_*IR*_, 5.7368 [mM/s] (yellow) and 0.01 [mM/s] (purple). For each case, we simulate until an apparent steady state for a range of *kh_y_d* and plot the curves of minima and maxima at equilibrium. In the oscillatory region, the minima and maxima diverge and show an oscillatory regime, denoted by yellow dashed line. Next, we hold k_hyd_ constant and vary V_*IR*_ to obtain the stability profiles for **c)** mTORC1 and **d)** mTORC2 dependent on V_*IR*_ For two values of *kh_y_d*, 1 × 10^-4^ [mM/s] and 0.149 [mM/s] we simulate until an apparent steady state for a range of V_*IR*_, then plot the curves of local minima and maxima at equilibrium for mTORC1 and mTORC2. For regions of monostability, outside of dashed lines, the curves of minima and maxima converge to the steady state. In oscillatory regimes, within the dashed lines, the local minima and maxima due to oscillations form an envelope. The size and shape of the envelope is dependent on both k_hyd_ and V_*IR*_. Then, to characterize this relation in a 2D parameter space, we plot corresponding 3D surface plots for **e)** mTORC1 and **f)** mTORC2.

**Figure 7:**
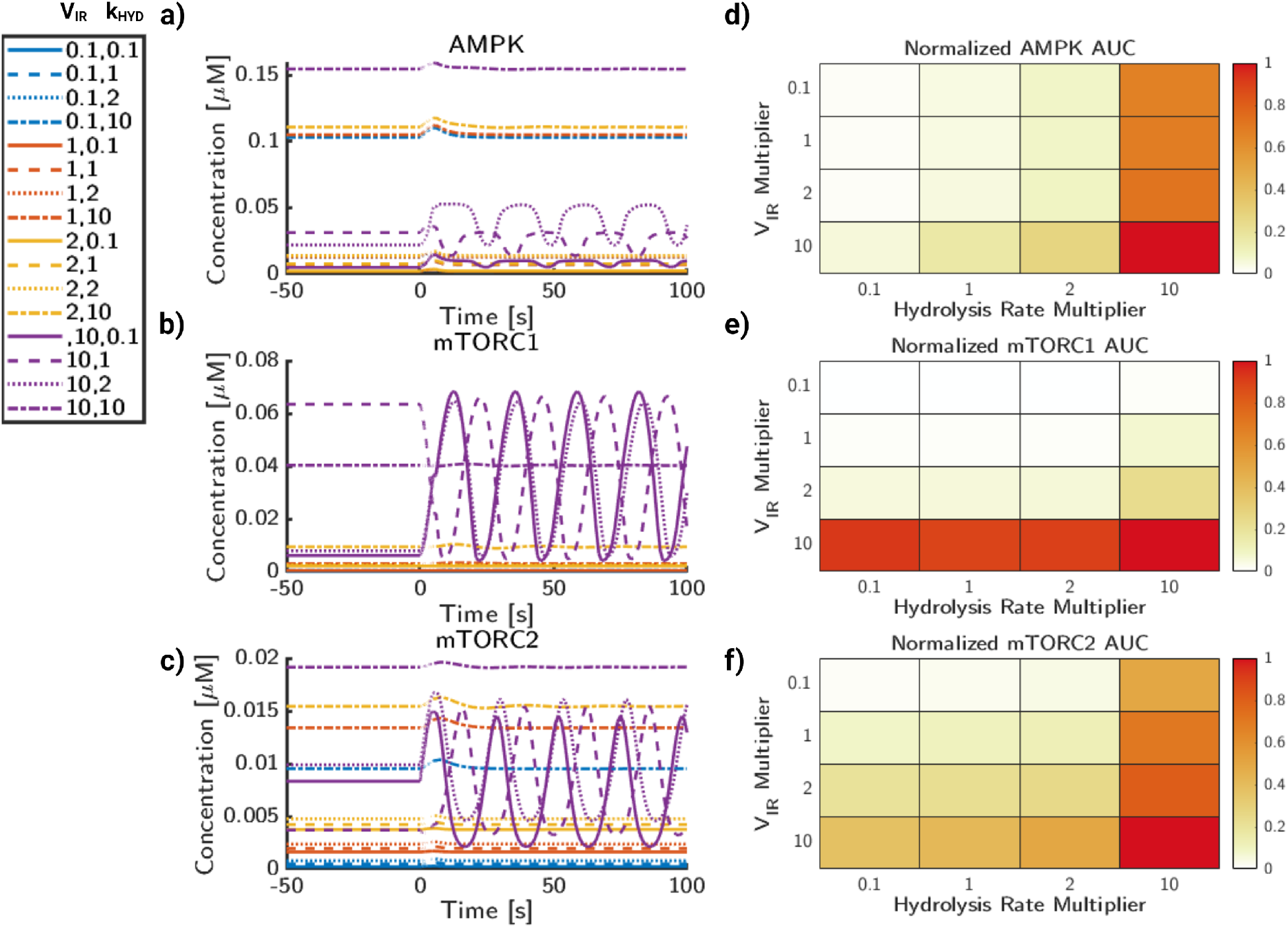
Insulin and cellular metabolism govern phosphorylation rates of AMPK, mTORC1, and mTORC2. **a)** AMPK phosphorylation concentration trajectories for a range of values for insulin receptor activity,V_*IR*_, and ATP consumption rates, *k_hyd_*. **b)** mTORC1 phosphorylation concentration trajectories for a range of values for insulin receptor activity, V_*IR*_, and ATP consumption rates, *k_hyd_*. **c)** mTORC2 phosphorylation concentration trajectories for a range of values for insulin receptor activity,V_*IR*_, and ATP consumption rates, *k_hyd_*. **d)** AMPK AUC normalized to the maximum AUC value of AMPK. **e)** mTORC1 AUC normalized to the maximum AUC value of mTORC1. **f)** mTORC2 AUC normalized to the maximum AUC value of mTORC1.

**Figure 8:**
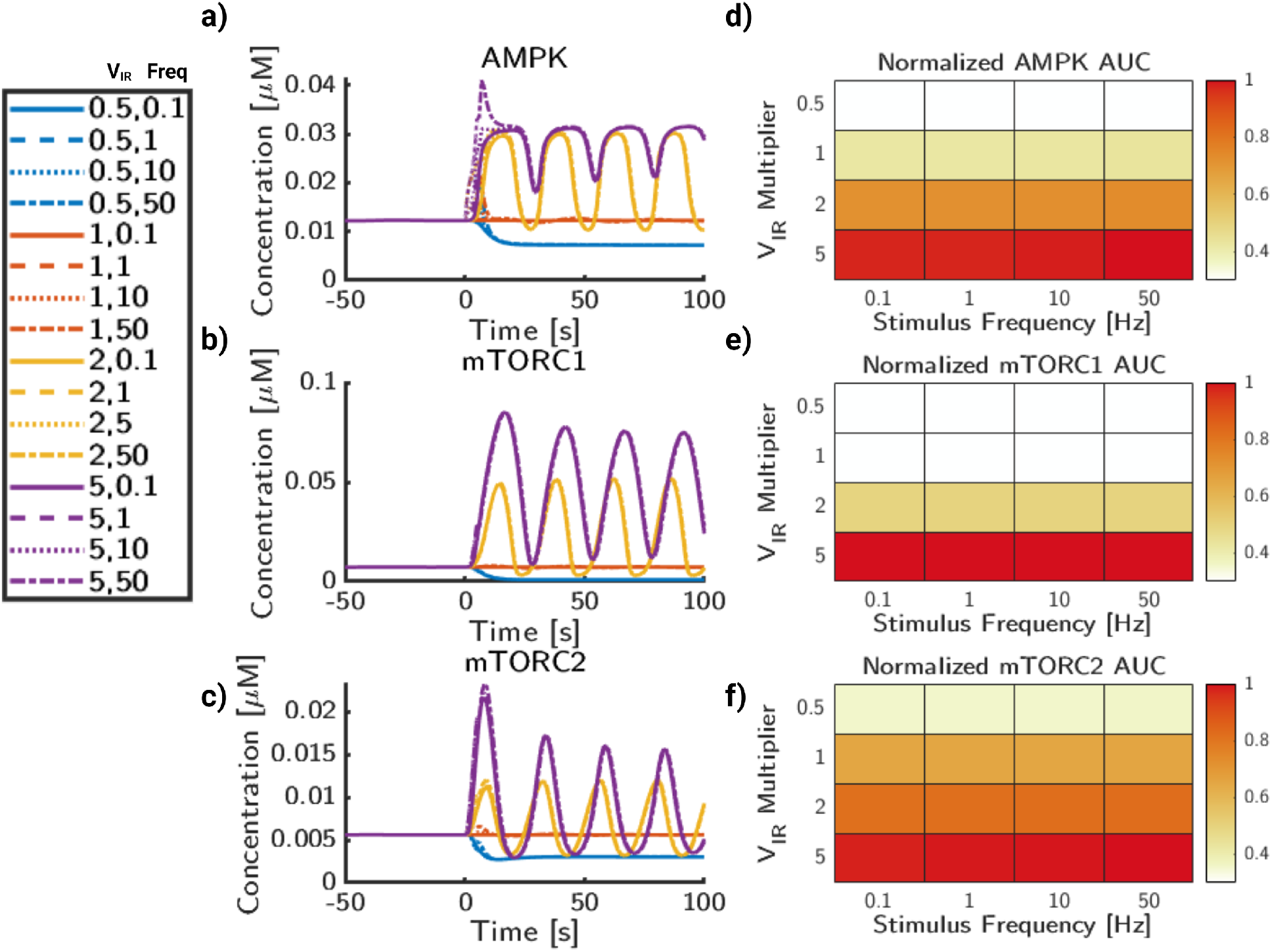
Insulin signaling and glutamate frequency influence dynamic behavior of AMPK and mTOR phosphorylation. **a)** AMPK phosphorylation concentration trajectories for a range of values for insulin receptor activity and glutamate stimulus frequency. **b)** mTORC1 phosphorylation concentration trajectories for a range of values for insulin receptor activity and glutamate stimulus frequency. **c)** mTORC2 phosphorylation concentration trajectories for a range of values for insulin receptor activity and glutamate stimulus frequency. **d)** AUC for AMPK normalized to the maximum AUC value of AMPK. **e)** AUC for mTORC2 normalized to the maximum AUC value of mTORC1. **f)** AUC for mTORC2 normalized to the maximum AUC value of mTORC2.

**Figure 9:**
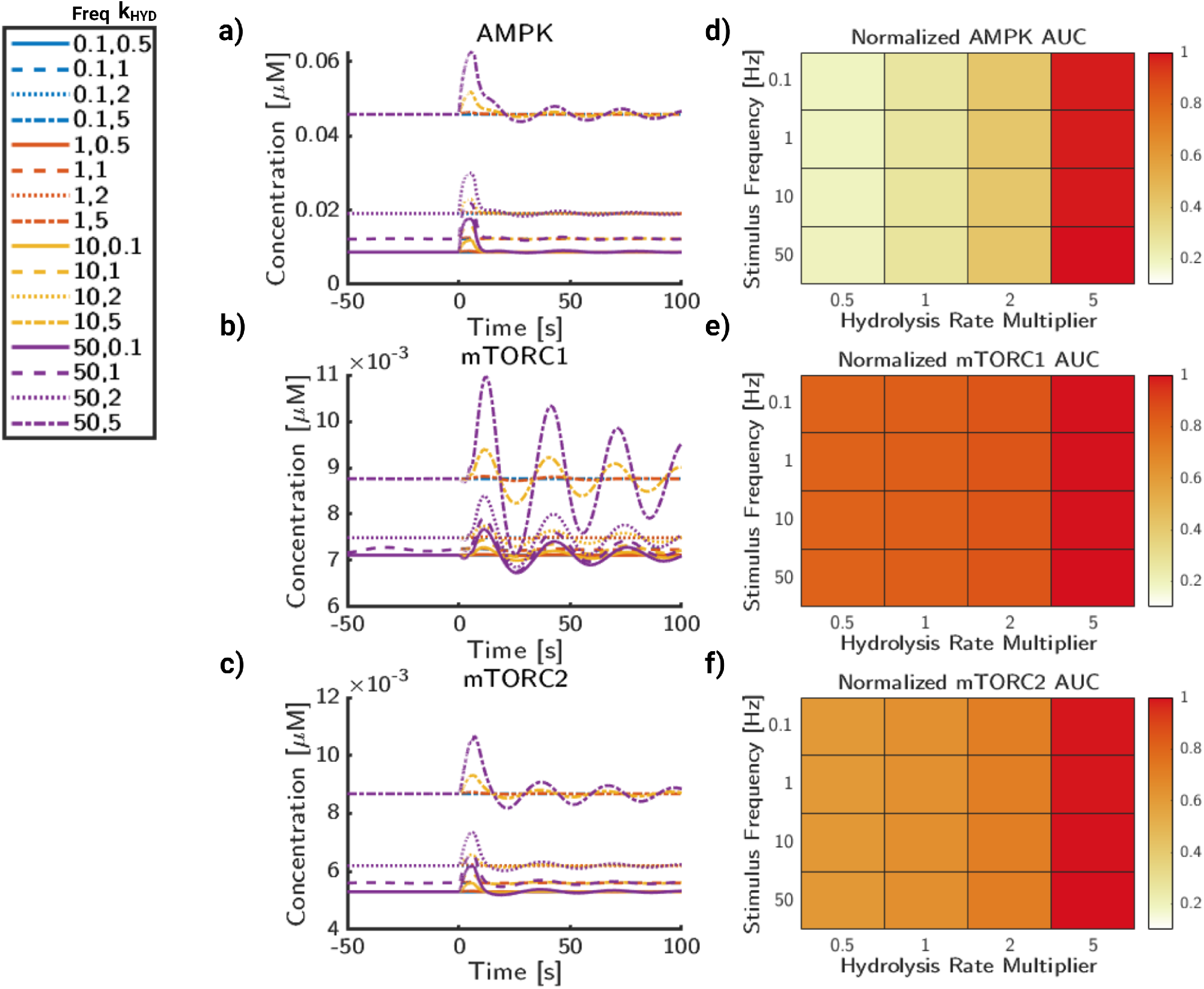
Glutamate frequency and metabolic stress lead to increases in AMPK and mTOR deviations. **a)** AMPK phosphorylation trajectories for a range of values for glutamate frequency and ATP consumption rates, *k_hyd_*. **b)** mTORC1 phosphorylation trajectories for a range of values for glutamate stimulus frequency and ATP consumption rates, *k_hyd_*. **c)** mTORC2 phosphorylation trajectories for a range of values for glutamate stimulus frequency and ATP consumption rates, *k_hyd_*. **d)** AUC for AMPK normalized to maximal AUC value. **e)** AUC for mTORC1 normalized to maximal AUC value. **f)** AUC for mTORC2 normalized to maximal AUC value.

Finally, we varied both *k_hyd_* and V_*IR*_ and plot the same curves as surface plots in 3D. For mTORC1 (Figure 6e), rising V_*IR*_ leads to an increase in the steady state. Furthermore, increasing *k_hyd_* reduces the range of values that produce oscillatory behavior. Outside the oscillatory regime, mTORC1 has a monotonic relationship with V_*IR*_. Within the parameter range studied, the steady state behavior of mTORC1 is strongly related to the magnitude of V_*IR*_, while *k_hyd_* has diminishing impact past the oscillatory envelope. For mTORC2 (Figure 6f), a similar regime of oscillations was found. However, for mTORC2, V_*IR*_ does not strongly affect the magnitude of mTORC2 at steady state. In contrast to mTORC1, in which V_*IR*_ was the primary controller of mTORC1 steady state, *k_hyd_* results in a much stronger increase mTORC2 steady state for all values of V_*IR*_. In summary, the oscillations of mTORC1 and mTORC2 that are observed in the model are dependent on the values of *k_hyd_* and V_*IR*_ and we predict that high energy consumption states (high values of *k_hyd_*) result in monostable steady states, while low values of *k_hyd_* and enable steady state oscillations within a range of V_*IR*_ values.

### Signaling frequency, ATP consumption, and insulin signaling modulate system response to energy stress

Thus far, we have investigated the impact of glutamate frequency alone (Figure 4) and the bifurcation behavior of the metabolic pathways in Figure 6. Next, we show glutamate stimulus frequency, IRS, and hydrolysis rate combine to influence the energy state of the system. In Figure 7, we observe the trends associated with changes in V_*IR*_ and *k_hyd_* under a 10 Hz stimulus pulse for 5 seconds. Trajectories for pAMPK, mTORC1, and mTORC2 are shown in Figure 7a-c. In these trajectories, the line color refers to the magnitude of V_*IR*_ while the line markers represent the *k_hyd_* magnitude. Qualitatively, increases in *k_hyd_* correspond to small concentration shifts of the trajectories predicted by V_*IR*_. For clarity, we will refer to the parameter values relative to the base model values (V_*IR,b*_ and *k_hyd,b_*). Concentration trajectories for AMPK are shown in Figure 7a. We observed that for most trajectories, the peak activity corresponds with the time of the stimulus. Additionally, for the case of V_*IR*_ = 10 × V_*IR,b*_ (purple lines) for AMPK, the rate of *k_hyd_* appears to affect the phase of the oscillations as well. Interestingly, this does not appear to be reflected in any other value of V_*IR*_ as the only oscillations of large magnitude at this frequency of stimulus appear to be due to V_*IR*_.

For mTORC1 (Figure 7b), we see a similar trend in the curves generated by increasing V_*IR*_ values to 10 × V_*IR,b*_. While many of the curves have a relatively low steady state and transient concentration as a result of the 10 Hz stimulus, The phase shift observed with AMPK in Figure 7a also leads to a shift in phase for mTORC1, however, the steady-state of these curves and direction of the oscillations shifts. This is exemplified most by 1 × *k_hyd,b_* and 2 × *k_hyd,b_*, in which the relative size magnitudes of oscillations are similar in magnitude, however the initial steady state is different. Furthermore, oscillations are abrogated by higher values of *k_hyd_*, which is consistent with the higher non-oscillatory regime of mTORC1 found in Figure 6e. For mTORC2 (Figure 7c), there are a spectrum of states available to mTORC2 phosphorylation dependent on *k_hyd_*, but more strongly on V_*IR*_ value. Similar to mTORC1, mTORC2 has a phase shift in the oscillations, but not necessarily the direction of the oscillations as the steady states are more similar between 0.1,1, and 2 × *k_hyd,b_*.

The relative AUC of the trajectories in Figure 7a-c are then quantified and summarized in Figure 7d-f. Since the curves can be vastly different in magnitude, for ease of comparability all AUC’s are ratios of the AUC of an unstimulated system under the same integration duration and then normalized to the maximum AUC for the selected trajectories. For AMPK (Figure 7d), increasing V_*IR*_ and *k_hyd_* both lead to increases in AUC, with larger increases correlated to higher magnitudes of *k_hyd_*. For mTORC1 (Figure 7e), increasing V_*IR*_ increases mTORC1 AUC, but increasing *k_hyd_* leads to slight increase in mTORC1 AUC. Finally, for mTORC2 (Figure 7f), the AUC for most of the trajectories are similar to AMPK. However, the trend follows that increasing both V_*IR*_ and *k_hyd_* also increases the mTORC2 AUC, and perhaps mTORC2 has the highest sensitivity to *k_hyd_*. The largest magnitude increase of mTORC2 activation comes from when both *k_hyd_* and V_*IR*_ are highest.

We next analyzed the interaction of the insulin substrate strength, V_*IR*_, with the glutamate stimulus frequency (Figure 8). V_*IR*_ influences the steady state and oscillatory values of AMPK activity (Figure 8a), which does not appear to be as strongly impacted by glutamate frequency on this concentration scale. But, as shown previously in Figure 4, higher frequencies above 10 Hz induce large transient pulses of AMPK activity. The primary influence of frequency appears to be increasing the activity of AMPK during stimulus. mTORC1 (Figure 8b) is significantly more sensitive to the magnitude of V_*IR*_ than to the input frequency. This is the case for the high-frequency cases of both 0.1 and 10 × V_*IR*_, in which no dampened oscillations are shown after approximately 50 seconds. A similar case is observed with mTORC2 (Figure 8c), in which both V_*IR*_ and stimulus frequency impact the initial peak height. However, particularly at cases of high frequency and also high V_*IR*_ magnitude, there are high amplitude oscillations and a delayed return to steady state. Oscillations are still observed in 1 and 2 × V_*IR*_ cases, but are less pronounced due to the wider range of magnitudes from higher frequencies.

The relative AUC (Figure 8d-f) is quantified in the same process as Figure 7. For AMPK (Figure 8d), the relative AUC increases more with VIR than stimulus frequency. The highest magnitude of AUC for AMPK comes from both high stimulus frequency as well as high V_*IR*_ magnitude. For mTORC1 (Figure 8e), while there is a clear increase in AUC while V_*IR*_ magnitude increases, any differences due to frequency are minimal. This implies that the stimulus frequency does not impact the overall signaling capacity of mTORC1, but may impact the dynamics of signaling on short timescales. Likewise, mTORC2 (Figure 8f) similarly shows a dependence on V_*IR*_ magnitude for overall mTORC2 AUC. However, as frequency increases, the AUC increases, but at a lower rate than changing the insulin receptor signaling strength.

Finally, we analyze interactions between signaling frequency and ATP consumption rate (Figure 9). As before, stimulus frequency appears to be the largest driver of transient stimulus for AMPK (Figure 9a). As frequency increases, the time AMPK returns back to equilibrium is delayed, especially for higher magnitudes of *k_hyd_*. The rate of *k_hyd_* can also significantly raise the baseline values of AMPK activation, which is consistent with its model function and AMPK’s activation conditions as the AMP/ATP ratio increases with higher *k_hyd_*. For mTORC1 (Figure 9b), *k_hyd_* does not have a strong impact to the activation rates of mTORC1, except at very high rates of *k_hyd_*. The stimulus frequency, however, has a strong correlation with increased amplitude of mTOR1. For mTORC2 (Figure 9c), frequency again shows an increased transient amplitude increase, but not a significant change to the overall trajectories. Likewise, *k_hyd_* only appears to significantly change the state of mTORC2 when it is very high, 10 × *k_hyd,b_*

The relative AUC (Figure 9d-f) is quantified in the same process as Figure 7. For AMPK (Figure 9d), the relative AMPK AUC shows a strong dependence on *k_hyd_*, but also a minor increase due to frequency. For mTORC1 (Figure 9e), frequency changes appear to affect the dynamics of the system, but not necessarily the AUC. While the initial peak heights increase, this does not appear to impact the overall signaling capacity of the system. However, this may lead to differences in short timescale activation profiles of mTOR and its downstream targets. For mTORC2 (Figure 9f), the AUC shows an overall increase with hydrolysis rate. It also displays a minor increase due to signaling frequency, particularly at higher hydrolysis rates. In summary, we have shown that crosstalk between glutamate frequency and metabolic signaling plays an important role in protein kinase activity, which has implications on cell fate and neuronal function. Glutamate signaling frequency appears to control the dynamic behavior, including the amplitude change and observed oscillations, however cellular energy consumption controls the steady state, which has a much stronger influence over AUC magnitude.

## Discussion

In this study, we have investigated the crosstalk between calcium influx in dendritic spines and mTOR and AMPK activation bridging multiple timescales of signaling in the context of dendritic spines using computational modeling. The pathways explored here have significant implications for synaptic plasticity as the downstream pathways of mTOR have been shown to be necessary for LTP and LTD [31]. Our main predictions focus on the possible divergent stability regimes in the system as a function of crosstalk between signaling and metabolic pathways. We find that the model exhibits a parameter-dependent oscillatory behavior, which can enable neurons to tune the cellular response to stimulus-based upon energy consumption, extracellular factors like insulin signaling, and signaling frequency.

It is believed that synaptic plasticity is triggered mainly through high-frequency signaling [57]. While the exact mechanisms contributing to synaptic plasticity are unknown, various works have shown that it is dependent on subcellular structure, calcium signaling, metabolics, and many other intermediary proteins including mTOR [4, 58, 63, 64]. Additionally, the mechanisms contributing to synaptic pruning and loss of dendritic spine density are unknown. In such complex processes, mathematical modeling may be able to contribute strongly to our under-standing of crosstalk between signaling, metabolism, and synaptic plasticity.

We predict that the cellular response to energy consumption and external cellular signaling are frequency dependent and produce ranges of cellular stability behavior. Specifically, the range of oscillations between low and high energy states can contribute to distinct signaling mechanisms. Furthermore, the oscillatory state may serve as a buffer range between random or noisy neuronal signals and high frequency signals that invoke LTP in synapses. While it is known that AMPK hyperactivation can contribute to synaptic pruning [65], these oscillations allow the system to reach higher rates of AMPK activation without raising the relative AUC. In this sense, oscillations controlled by the insulin system may regulate the levels of energetic stress that a neuron can withstand before triggering autophagy pathways. Although technically challenging, pharmacological experiments for AMPK and mTOR activators in dendritic spine could show a change in sensitivity to glutamatergic stimulus and subsequently affect LTP and LTD.

Models of synaptic plasticity can be generalized to phenomenological models (including rate based models and spike timing based models) and biophysical models which generally focus on calcium signaling and downstream signal transduction via CAMKII [7,66,67]. However, looking forward, models of synaptic plasticity can incorporate crosstalk effects of insulin on downstream components in biophysical models of synaptic plasticity. Insulin signaling has been shown to regulate neuroplasticity in both developing and adult brains [68]. While this can be explained by reports that insulin can transiently induce potentials via NMDA [69, 70], insulin has also been shown to impair NMDA dependent LTP [71]. Similarly, insulin may also attenuate AMPAR signaling by inducing AMPA receptor internalization [72]. However, the mechanism of action for these disparate actions of insulin needs to be understood in the context of the dynamics of intracellular signaling molecules like AMPK and mTORC.

AMPK may be critical in the low-energy response of neurons, both from an intraceullar and systems level perspective [27,73]. While AMPK is critical for the integration of inputs, mTORC is linked to protein translation and autophagy, which can serve as outputs for influencing cellular state. mTORC1, while active in presynaptic neurons, is not expressed as heavily in postsynaptic dendritic spines [31]. mTORC2, is expressed in dendritic spines and activated during synaptic signaling [31,73,74]. There are additional reports that mTORC2 is essential for synaptic plasticity [31,75]. Due to the importance of insulin signaling and AMPK, further investigation of the kinetics of mTOR is vital for our understanding of cellular interactions leading to synaptic plasticity.

Future works in both modeling and experiments may focus more on building upon knowledge of synaptic plasticity. For modeling complex biochemical pathways, deterministic solver methods are feasible due to high molecule numbers [76]. However, incorporating stochastic solver methods may be able to capture biological behavior, which is particularly important in multi-scale problems like dendritic spines [3, 77] Finally, multivariate experimental measurements in spines during synaptic activity would be need to constrain the models and test model predictions.

## Abbreviations

AMP: Adenosine Monophosphate
ADP: Adenosine Diphosphate
ATP: Adenosine Triphosphate
AMPK: AMP-activated protein kinase
mTORC1: Mammalian Target of Rapamycin Complex 1
mTORC2: Mammalian Target of Rapamycin Complex 2
AKT: Protein Kinase B
ULK1: UNC-51 like autophagy activating kinase 1
AMPAR: α-amino-3-hydroxy-5-methyl-4-isoxazolepropionic acid receptor
NMDAR: N-methyl-D-aspartate receptor
SIRT1: Sirtuin 1
GLUT1-4: Glucose Transporter 1-4
IP3: Inositol 1,4,5-trisphosphate
IP3R: IP_3_ receptor
RyR: Ryanodine receptor
SERCA: sarcoplasmic/endoplasmic reticulum calcium ATPase
ER: Endoplasmic reticulum
PMCA: Plasma Membrane Calcium ATPase
NCX: Na^+^/Ca^2+^ exchanger
mGluR: Metabotropic glutamate receptor
PKC: Protein kinase C
PIP2: Phosphatidylinositol (4,5)-bisphosphate
LTP: Long-Term Potentiation
LTD: Long-Term Depression
CAMKK2: Calcium-Calmodulin Kinase Kinase 2
LKB1: Liver Kinase B1
PI3K: Phosphoinositide 3-kinase

## Data Availability Statement

All code and data needed to reproduce this work are included in public repository hosted at: https://github.com/aleung15/AMPKmTOR2022.

## Acknowledgments

The authors would like to acknowledge Maria Hernández Mesa, Guadalupe Garcia, and Lingxia Qiao for their feedback and discussion. Additionally, we acknowledge biorender.com for tools used to generate graphics in Figure 1. Finally, this work was supported by the University of California, San Diego Interfaces Graduate Training Program, Air Force Office of Scientific Research (AFOSR) Multidisciplinary University Research Initiative (MURI) grant FA9550-18-10051.

## Supplementary Figures

**Supplemental Figure 1:**
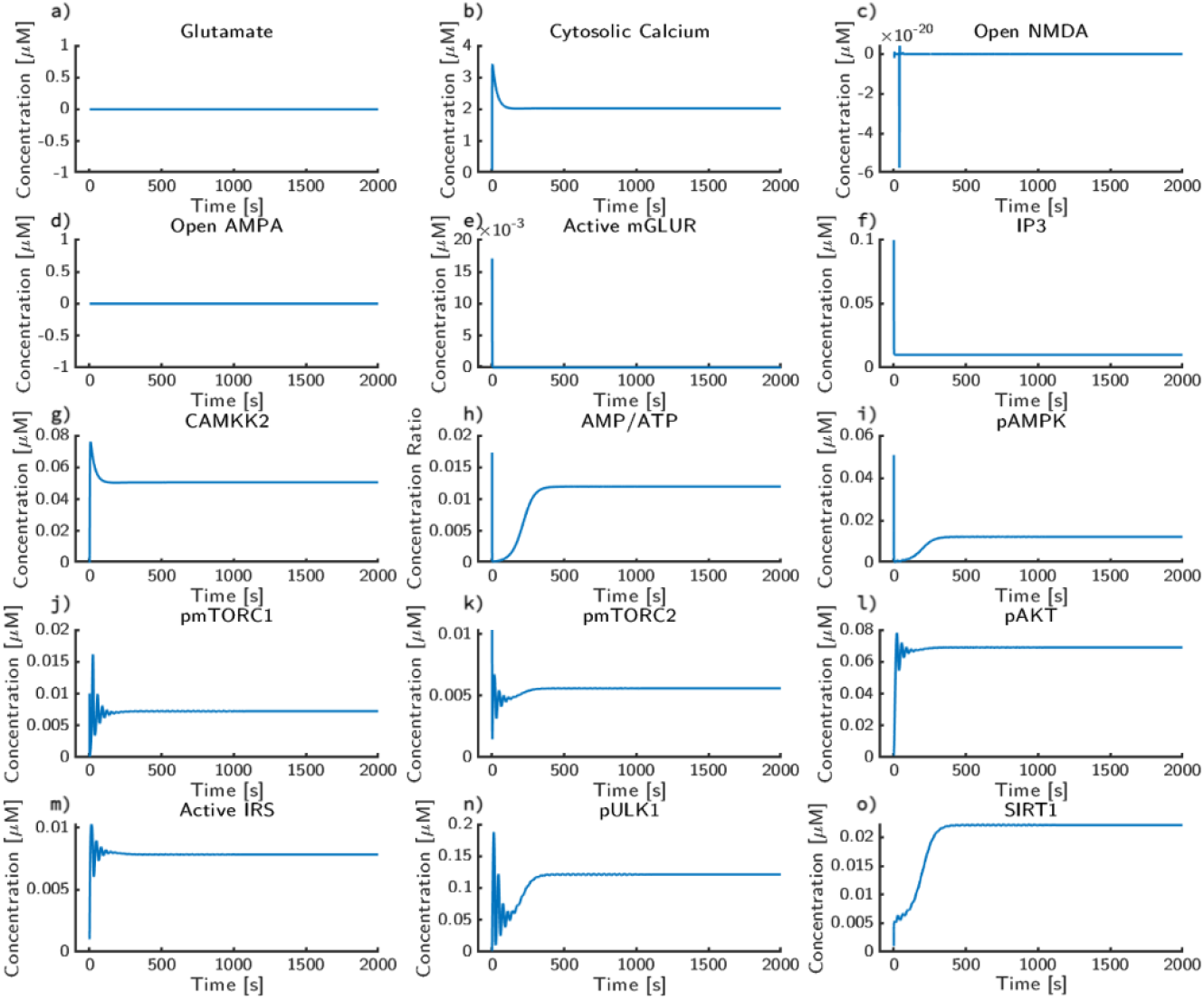
Equilibration simulation for system without glutamatergic stimulus. The system re-cieved a single pulse of glutamate and was allowed to equilibrate for 2000 seconds. Trajectories are plotted for:**a)** glutamate, **b)** cytosolic calcium, **c)** open NMDA Receptors, **d)** open AMPA receptors, **e)** active mGLUR receptor, **f)** IP3, **g)** CAMKK2, **h)** AMP/ATP ratio, **i)** phosphorylated AMPK, **j)** phosphorylated mTORC1, **k)** phosphorylated mTORC2, **l)** phosphorylated AKT, **m)** active IRS, **n)** phosphorylated ULK1, **o)** SIRT1.

